# Cyp26b1 restrains murine heart valve growth during development

**DOI:** 10.1101/2021.07.05.450958

**Authors:** Neha Ahuja, Max S. Hiltabidle, Hariprem Rajasekhar, Haley R. Barlow, Edward Daniel, Sophie Voss, Ondine Cleaver, Caitlin Maynard

## Abstract

Endothelial cells (ECs) are critical to proper heart valve development, directly contributing to the mesenchyme of the cardiac cushions, which progressively transform into mature valves. To date, investigators have lacked useful markers of valve ECs to fully evaluate their contributions during valve morphogenesis. As a result, it has been unclear whether the well-characterized regional differentiation of valves correlates with any endothelial domains in the heart. Furthermore, it has been difficult to ascertain whether endothelial heterogeneity in the heart influences underlying mesenchymal zones in an angiocrine manner. To identify regionally expressed EC genes in the heart valves, we screened publicly available databases and assembled a toolkit of endothelial-enriched genes. We identified Cyp26b1 as one of many endothelial enriched genes found to be expressed in the endocardium of the developing cushions and valves. Here, we show that Cyp26b1 is required for normal heart valve development. Genetic ablation of Cyp26b1 in mouse embryos leads to abnormally thickened aortic valve leaflets, which is due in part to increased endothelial and mesenchymal cell proliferation in the remodeling valves. In addition, Cyp26b1 mutant hearts display ventricular septal defects (VSDs) in a portion of null embryos. We show that loss of Cyp26b1 results in upregulation of retinoic acid (RA) target genes, supporting the observation that Cyp26b1 has RA-dependent roles. Together, this work identifies a novel role for Cyp26b1 in heart valve morphogenesis. Understanding the spatiotemporal expression dynamics of cardiac EC genes will likely prove useful to the investigation of both normal as well as dysfunctional heart valve development.

**HIGHLIGHTS:**

· A mouse heart valve gene expression atlas can be generated with publicly available online tools, such as Genepaint and other gene expression databases.
· Endothelium of developing mouse heart valves is regionally heterogeneous.
· Cyp26b1 is expressed in the endocardial/endothelial lining of developing heart valves.
· Loss of Cyp26b1 leads to significant enlargement of aortic valves and to ventricular septal defects.
· Cyp26b1 represses cell proliferation in valve mesenchyme.
· Retinoic acid targets are upregulated in *Cyp26b1-/-* heart valves, indicating dysregulation of RA metabolism.

## INTRODUCTION

One percent of newborn infants suffer from congenital heart disease (CHD) (Virani et al., 2020) caused by cardiac developmental defects like valve malformations and VSDs (Hoffman and Kaplan, 2002; Lincoln and Yutzey, 2011). Mammalian hearts have four valves: two atrioventricular (AV) valves and two semilunar (SL) valves. Studies of valve development in embryonic mice have shown that at embryonic day 10.5 (E10.5), endothelial cells (ECs), also called endocardial cells in the heart, comprise a monolayer that encases the cushions and lines the heart. These ECs undergo a transition to mesenchymal tissue (an endothelial-to-mesenchymal transition called EndoMT), invading the endocardial cushion interstitium (Armstrong and Bischoff, 2004). By E14.5, the valves progressively take shape, as mesenchymal cells within the valve secrete extracellular matrix (ECM) proteins to give rise to different functional zones (Hinton and Yutzey, 2011). The uppermost zone of the valve (atrialis (AV)/ventricularis (SL)) experiences blood flow and is composed of flexible elastin-containing mesenchymal tissue. The central zone (spongiosa) contains loose collagen and proteoglycans, and the underlying zone (fibrosa) is composed of load-bearing collagen (Combs and Yutzey, 2009). These mesenchymal zones are lined by ECs, which serve to guide valvular development by integrating signals from myocardial cells and blood flow.(Tao et al., 2012). The interaction between overlying endothelial domains and the underlying mesenchymal zones at later stages bears further investigation.

Notably, valve endothelium is not uniform. During valve development, ECs that overlie the atrialis/ventricularis (inflow endothelium) experience increasingly higher shear stress than ECs that overlie the fibrosa (outflow endothelium), leading to functional and molecular differences between these endothelial populations (Butcher and Nerem, 2007; Chester and Taylor, 2007). These differences are perhaps best illustrated in the context of aortic valve calcification, a disease in which calcium deposits develop exclusively on the outflow side of the aortic valve (El-Hamamsy et al., 2009). The inflow aortic valve endothelium secretes Nitric Oxide, which is thought to repress the development of calcific nodules (Richards et al., 2013). Indeed, transcriptomic profiling of adult porcine valve leaflets identified over 585 differentially expressed genes between the outflow and inflow endothelium (Simmons et al., 2005). Recent advances in single cell RNA-sequencing have enabled the identification of three distinct valve EC populations present in aortic and mitral murine valves (Hulin et al., 2019). Similarly, valvular cells in the fetal human heart show distinct subtypes in SL and AV valves (Cui et al., 2019). How this endothelial heterogeneity arises and functionally contributes to valve development is not well understood.

To investigate the molecular basis for valve endothelial heterogeneity, we utilized publicly available data sets to screen for genes with regionally restricted expression patterns during valve development. We identified several novel genes expressed in the valve endothelium and selected Cyp26b1 for further investigation. Cyp26b1 is part of the cytochrome P450 family of enzymes and metabolizes retinoic acid (RA) to inactive forms. RA directs key steps in cardiac development, including cardiac specification, looping, and endocardial cushion formation [reviewed in (Pan and Baker, 2007)]. Exogenous administration of RA in avian embryos at Hamburger-Hamilton (HH) stage 34 results in VSDs (Broekhuizen et al., 1995), while treatment at an earlier stage resulted in less severe defects, including restricted AV cushion volume and changed hemodynamics (Bouman et al., 1998). In murine models, intraperitoneal injection of all-trans RA into pregnant dams at E8.5 resulted in decreased EndoMT in the outflow cushion by E10.5 (Sakabe et al., 2012). Mutation of *Raldh2* – an RA synthesizing enzyme – resulted in excess mesenchyme in the outflow tract by E13.5 (El Robrini et al., 2016). Together, these results highlight the essential role that RA signaling plays during valve development with further emphasis on regional effect.

The Cyp26 family of enzymes has been shown to have essential roles during development of several organ systems. Mouse *Cyp26b1* mutants display shortened limbs due to decreased chondrogenesis and consequent defects in skeletal formation (Dranse et al., 2011). We previously found that during murine lung development, *Cyp26b1* mutants show defects in differentiation of alveolar type1 (AT1) cells, as well as failures of epithelial morphogenesis (Daniel et al., 2020). In zebrafish, depletion of Cyp26a1 and Cyp26c1 leads to hypocellular outflow tracts (OFTs) due to a deficit of second heart field progenitor migration (Rydeen and Waxman, 2016). These data show that the Cyp26 family of enzymes are key mediators of organogenesis; however, the role of Cyp26b1 in cardiac valve remodeling has not previously been examined.

In this report, transcriptional analysis of publicly available databases allowed us to identify numerous genes expressed in different regions of the cardiac valve endothelium. Among these genes, we find Cyp26b1 to be uniquely restricted to the endocardium of murine heart valves and expressed throughout valve formation. Furthermore, we find that *Cyp26b1-/-* hearts exhibit enlarged aortic valve leaflets secondary to excess outflow tract mesenchymal cell proliferation, as well as VSDs (the latter in approximately 30% of mutant hearts). We show that Cyp26b1 is required to limit proliferation during aortic valve development, to facilitate remodeling of the aortic valve leaflet, and to drive proper cardiac chamber septation. In addition, we show that loss of Cyp26b1 results in increased expression of RA target genes, suggesting increased RA signaling and underscoring an important role for RA in normal heart development. Together, our findings provide a road map for identification of novel heart EC genes and demonstrate a previously unknown role for the Cyp26b1 enzyme during heart development.

## RESULTS

### Identification of genes enriched in valve endothelium

The endothelium of vertebrate organs and tissues has long been demonstrated to be highly heterogeneous, however the origin of this heterogeneity remains poorly understood. In addition, endothelium has been shown to participate in active crosstalk with the tissues it pervades. Examples of such angiocrine signals have steadily been identified over the last few decades (Ding et al., 2011; Lammert et al., 2001; Matsumoto et al., 2001; Rafii et al., 2016). To begin to understand EC signals, we must first fully characterize the dynamics of EC gene expression.

We set out to identify genes regionally enriched in the endothelium of the developing heart. Using the publicly available database Genepaint (https://gp3.mpg.de/), we examined images of RNA in situ hybridization (ISH) for over 2000 genes to assess valve specific expression patterns in E14.5 mouse embryos (**Fig. 1A**). This was done via a ‘stringent expression pattern’ search parameter on the website for heart structure with strong expression of regional patterns, followed by progressively more tolerant search criteria. The results yielded a variety of genes displaying various valve expression patterns, most of which were also expressed in ECs across other embryonic tissues, as well as other cell types. We focused on sections through heart valves, when available in the datasets (AV and SL valves, **Fig. 1B**). Within the heart, patterns were sometimes restricted to the endothelium of the valves or the underlying developing valve mesenchyme, or both. In some cases, the genes were also expressed in the endocardium.

**Figure 1.**
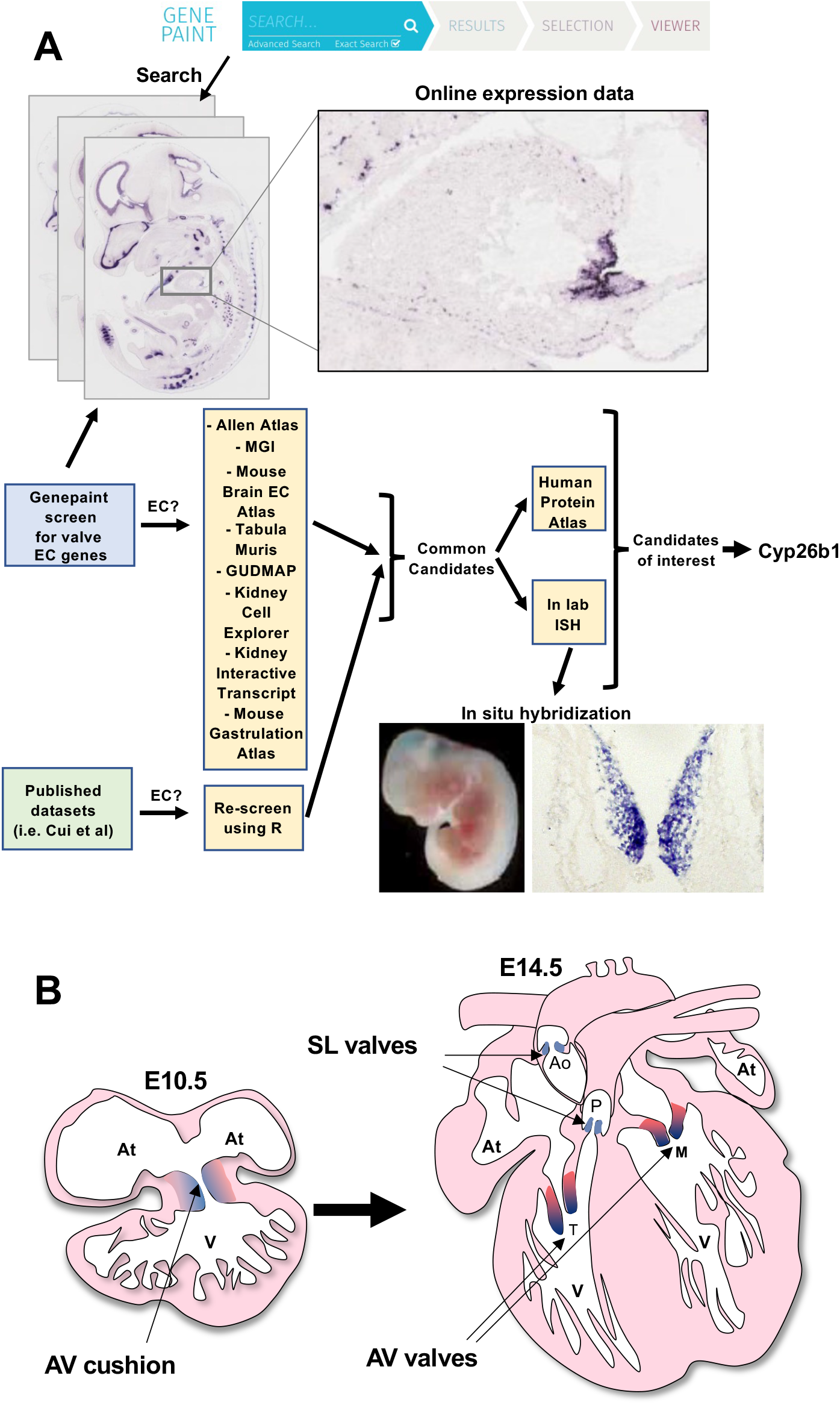
Screen for genes expressed in developing heart valves. (**A**) To identify regionally expressed endothelial genes expressed in the early heart, we screened publicly available databases (listed in text of manuscript). Using these tools, we examined *in silico* over 2000 genes that exhibited expression in the endothelium of large and small vessels, as well as in the endocardium. The Genepaint database is limited to expression data at embryonic day 14.5 (E14.5), although some earlier or later expression patterns could be found in the Allen Brain Atlas and Jax labs databases. Images of the heart were cropped and catalogued (see **Fig. S1**). We examined those genes that were visibly expressed by ISH and ssRNAseq analyses in either the tricuspid, atrioventricular, aortic or pulmonary valves of the embryonic heart. (**B**) Diagram of early heart cushions (E10.5) and developing heart valves (E14.5) that were considered in this analysis.

Our screen identified around 150 genes that exhibited regionalized expression patterns in different regions of the E14.5 heart valves, as per ISH assays available on Genepaint. This provided an initial picture of the wide variety of genes expressed across the heart valves at this midgestation stage. Endothelial-restricted gene expression of interesting candidates was further verified by assessing publicly available databases. We assessed whether candidate genes were expressed in endothelium of various tissues, such as the adult mouse brain (https://betsholtzlab.org/VascularSingleCells/database.html?gene=), the mouse heart and multiple adult mouse tissues (Tabula Muris, https://tabula-muris.ds.czbiohub.org/, Allen Brain Atlas (https://mouse.brain-map.org/, The Jackson Labs MGI Gene Expression Database (http://www.informatics.jax.org/gxd) the kidney (Gudmap https://www.gudmap.org/, Kidney Interactive Transcriptomics http://humphreyslab.com/SingleCell/displaycharts.php, Kidney Cell Explorer https://cello.shinyapps.io/kidneycellexplorer/), and the early developing embryonic primary vascular plexus (https://marionilab.cruk.cam.ac.uk/MouseGastrulation2018/) (see **Table 2**). In **Fig. S1**, we show gene expression patterns identified as endothelial in the databases listed above, underscoring EC heterogeneity in the developing mouse heart and cardiac valves (also see **Table 1**).

We also evaluated findings from other reports of transcriptional profiling of heart valves (Hulin et al., 2019; Cui et al., 2019). In one report, transcriptomic analysis of isolated fetal human hearts and human heart valves was carried out by the Tang group using single cell sequencing (Cui et al., 2019). We used this data set to find additional valve specific endothelial genes, and then cross-reference these genes against candidates from our Genepaint screen. We reasoned that genes on both lists would likely have roles in both mouse and human heart valve development. We performed unsupervised clustering analysis of the Tang data using the R package Seurat. We were able to identify 16 distinct clusters (**Fig. 2A**). To determine the identity of each cell cluster, we utilized expression of well-known marker genes. We first identified the endothelial cluster through expression of Pecam1, a well characterized endothelial gene (Privratsky et al, 2014) (**Fig. 2A,B**). We identified valve endothelium by assaying expression of Nfatc1, Klf2, and Ntrk2 (**Fig. 2C,D**), other known markers of the valve endothelium (Cui et al., 2019; Goddard et al., 2017; Wu et al., 2013). For markers used to identify other clusters, see **Fig. S2**.

**Figure 2.**
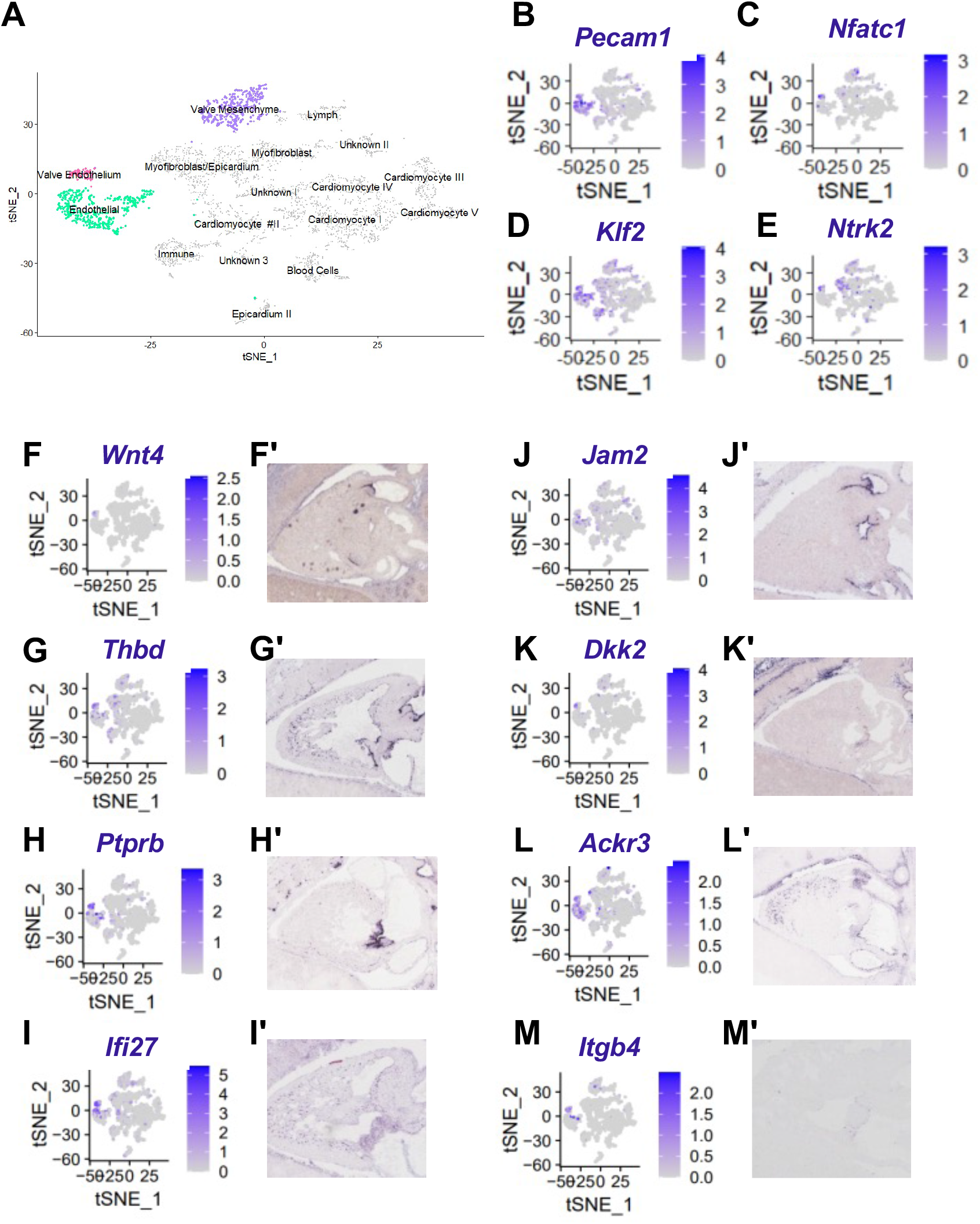
Gene expression in E14.5 mouse heart valves. (**A**) tSNE plot of human fetal cardiac cells from reclustering of the Tang group’s data. (**B**) tSNE of endothelial marker *Pecam1.* (**C-E**) tSNEs of valve endothelial markers *Nfatc1, Klf2, Ntrk1.* (**F**) tSNE and (**F’**) Genepaint ISH of known valve marker *Wnt4.* (**G-M**) tSNE and (**G’-M’**) Genepaint ISH of high-confidence valve specific genes *Thbd, Ptprb, Ifi27, Jam2, Dkk2, Ackr3,* and *Itgb4.*

To identify high confidence valve endothelial genes, we compared candidates from the Genepaint screen to the top differentially expressed genes in the valve endothelium from the Tang data. We identified several candidates expressed in both data sets (**Fig. 2F-M**). We also noted that many known markers of the valve endothelium (such as *Wnt4*) were identified by our screen (**Fig. 2F,F’**), thereby confirming the reliability of our approach. Several other novel markers of valvular ECs, including *Thbd, Ptprb, Ifi27, Jam2, Dkk2, Ackr3,* and *Itgb4,* were identified as well (**Fig. 2GM**). EC-specific expression of these genes in human hearts was confirmed by comparing EC enrichment to reported endothelial genes in human EC single cell sequencing data (The Human Protein Atlas, Single Cell Type Atlas www.proteinatlas.org). Together, this approach has yielded several high-confidence valve endothelial genes, that are ready for further experimental follow-up.

### Dynamic transcriptional heterogeneity of the developing valve endothelium

For approximately twenty of the most promising genes identified from either the Tang group’s sequencing data or our Genepaint screen, we generated Dig-labelled RNA probes and performed our own section ISH for examination at higher resolution and across additional stages. We first examined genes known to be expressed in the developing valves, including the endothelium. We were able to detect expression of these genes at E10.5 in the cushions and in the AV and SL valves at E14.5 (*Sox9, Wnt4, Car3* and *Ccn1* shown in **Fig. S3**). We were particularly interested in finding genes that displayed restricted expression in the valvular endothelium. We classified the endothelium into inflow, tip, and outflow domains, based on the direction that blood flows along the structure. We categorized the mesenchyme at E10.5 to proximal or distal domains in relation to the myocardial wall. At E14.5, we classified the mesenchyme based on the anatomical layers present within the leaflet: the fibrosa, spongiosa, and atrialis (AV) or ventricularis (SL) (**Fig. 3B**).

**Figure 3.**
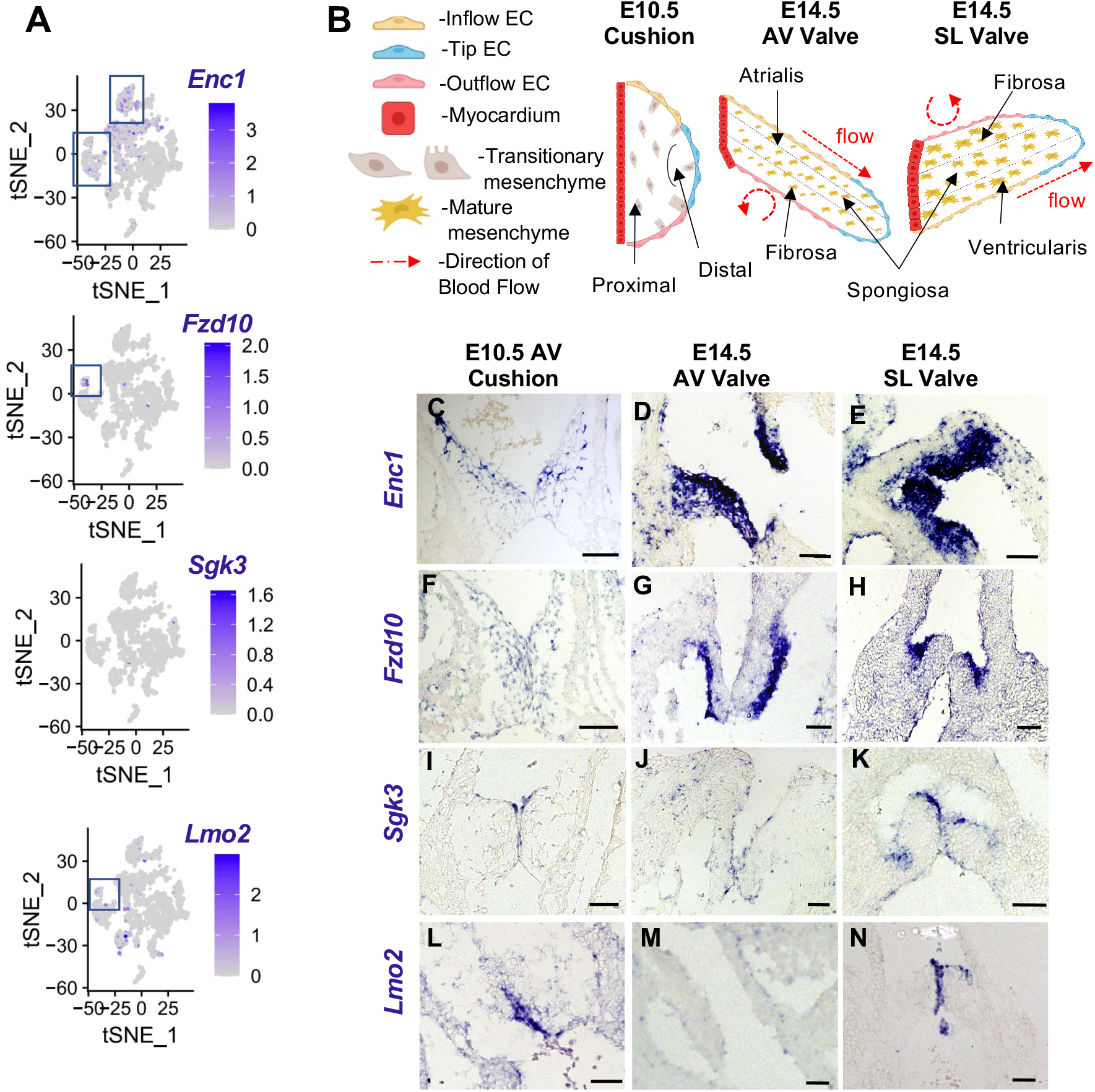
Regionalized expression of novel endothelial AV heart valve genes at E10.5 and E14.5. (**A**) t-SNE plots of *Enc1, Fzd10, Sgk3,* and *Lmo2.* (**B**) Diagrams of E10.5 cushions and E14.5 leaflets. Endothelial regions are defined as inflow, tip, and outflow. E10.5 mesenchymal regions are described as distal and proximal to the myocardial wall. E14.5 mesenchymal regions are fibrosa, spongiosa, and atrialis(AV)/ventricularis(SL). (**C-E**) ISH of *Enc1* in E10.5 and E14.5 valves. (**F-H**) ISH of *Fzd10* in E10.5 and E14.5 valves. (**I-K**) ISH of *Sgk3* in E10.5 and E14.5 valves.(**L-N**) ISH of *Lmo2* in E10.5 and E14.5 valves.

We next sought to identify endothelial genes expressed in the valves, which were not previously known to be involved in heart development. In **Fig. 3**, we show four genes, *Enc1, Fzd10, Sgk3,* and *Lmo2,* that demarcate different valve domains and have uncharacterized roles in valve development. We performed ISH of these genes at E10.5 and E14.5 and evaluated expression of these genes using t-SNE plots using the Tang group’s data (**Fig. 3A, C-N**). Enc1 is expressed specifically in the endocardial cushion outflow domain at E10.5; but by E14.5, Enc1 is expressed in both the endothelium and the valve mesenchyme in both the AV and SL valve leaflets (**Fig. 3C-E**). In agreement with this data, the t-SNE plot from the Tang group’s data suggests that *Enc1* is expressed strongly in the valve mesenchyme and endothelium of fetal human heart valves (**Fig. 3A**). At E10.5, Fzd10 is expressed at low levels but throughout the valve endothelium and mesenchyme (**Fig. 3F**). At E14.5 however, it becomes highly enriched in the outflow side of the valve, in both the AV and SL valves. Both endothelial and outer mesenchymal cells express *Fzd10* (**Fig. 3G,H**). At this later stage, no expression is detected in inflow endothelium or deep valve mesenchyme. *Sgk3,* by contrast, displays endothelium-specific expression in the valves, at both E10.5 and E14.5 (**Fig. 3I-K**). At both stages, expression in the endothelium is strongest along the inflow valve surfaces. At E14.5, *Sgk3* expression is notably stronger in the SL valve than in the AV valve. *Sgk3* was not detected in the Cui et al single cell RNA-sequencing data set (Cui et al., 2019)(**Fig. 3A**), potentially due to mouse-human difference in expression levels or to the small number of cells that express *Sgk3,* underscoring the importance of validation (as shown by our ISH). *Lmo2* by contrast was detected in the tip domain of the AV cushion at E10.5, expanded expression to both the inflow and outflow domains of the AV and SL valves by E14.5 (**Fig 3. L-N**). *Lmo2* also shows strong expression in the valve endothelial cluster by t-SNE (**Fig 3. A**). Together, these data highlight the EC heterogeneity of the developing valves and identify unique spatiotemporally dynamic patterns of gene expression in each valve.

### *Cyp26b1* is expressed in AV and SL valve leaflet endothelium throughout development

To investigate the role that a regionally restricted endothelial gene might play in valve development, we chose to further evaluate Cyp26b1, a retinoic acid metabolizing enzyme. Though Cyp26b1 was not robustly detected in the Tang group’s dataset (**Fig. S4A-A’**), it showed tightly restricted valve endothelial expression on Genepaint (**Fig. S1B).** We first sought to elucidate *Cyp26b1* expression pattern throughout cardiac valve development using RNA ISH. At E8.75, we observed branchial arch and intersomitic vessel expression, as previously reported (Bowles et al., 2014; MacLean et al., 2001), but no detectable expression in the heart (**Fig. S4A** and data not shown). At E9.5, we found that Cyp26b1 begins to be expressed in the cardiac endothelium (**Fig. 4A,A’**). By E10.5, *Cyp26b1* is expressed in the tip cushion endothelium of both the AV and outflow cushions (**Fig. 4B,C**). At E12.5, *Cyp26b1* is expressed on both the inflow and outflow domains of the SL valve endothelium and along the tip endothelium of the AV valves (**Fig. 4D-G**). At E14.5, *Cyp26b1* is expressed in both the inflow and outflow domains of the SL valve endothelium but becomes more restricted to the tip regions of the AV valves (**Fig. 4H-K**). By 18.5, *Cyp26b1* is expressed in the outflow endothelium of the SL valve leaflets and the tip endothelium of the mitral and tricuspid valve leaflets (**Fig. 4L-O**). Expression is strongest in the SL valve leaflets. Together, these data indicate that *Cyp26b1* is first expressed in both the developing AV and SL valves but is more strongly maintained in the SL valves.

**Figure 4.**
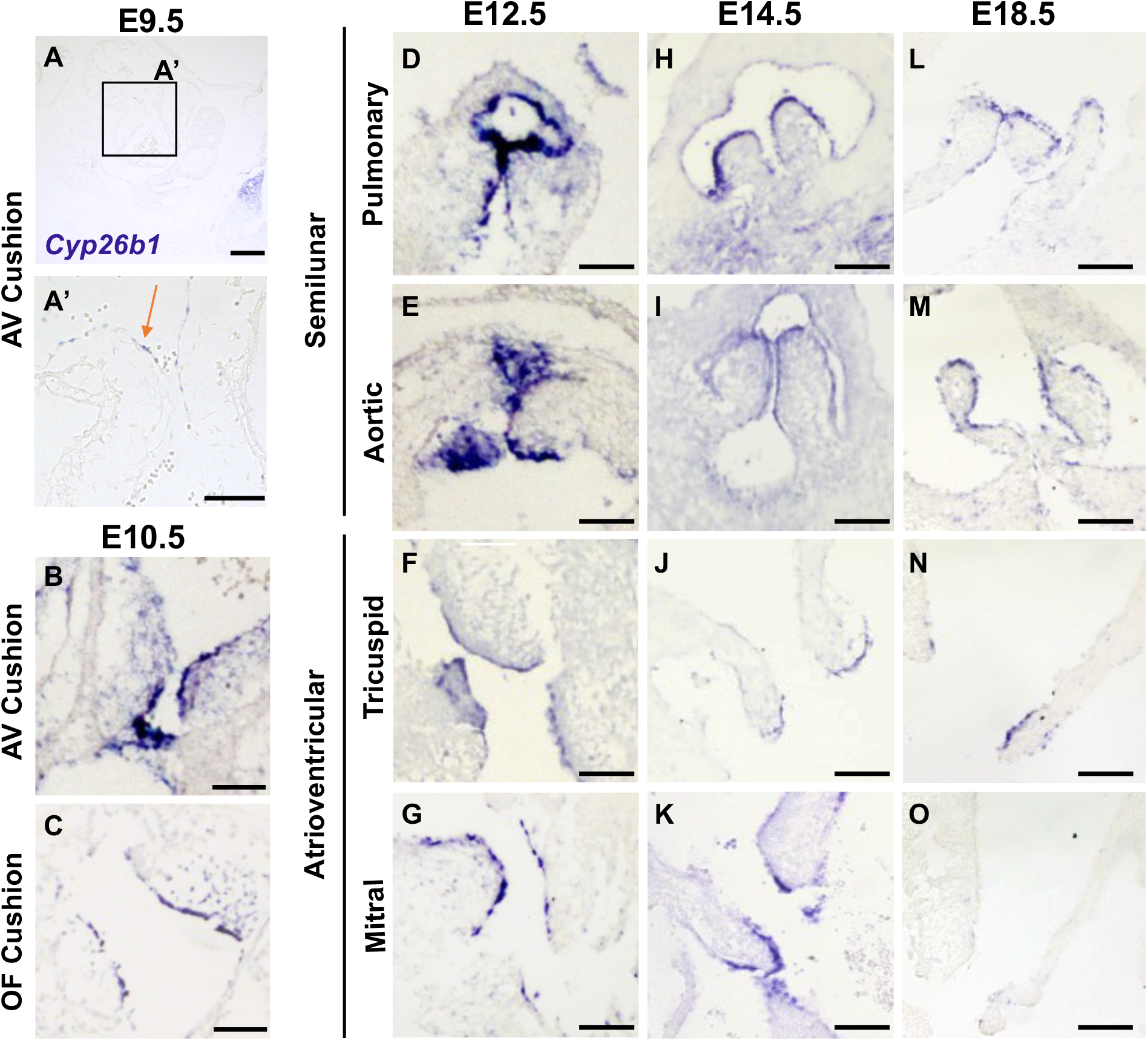
*Cyp26b1* expression is restricted to the valve endothelium throughout development. ISH of *Cyp26b1* in the (**A**) AV and (**B**) Outflow cushions at E10.5 (expression, red arrow). (**C-D**) ISH of *Cyp26b1* in the pulmonary valve between 12.5 and 18.5. (**E-F**) ISH of *Cyp26b1* in the aortic valve between 12.5 and 18.5. (**G-I**) ISH of *Cyp26b1* in the tricuspid valve between 12.5 and 18.5. (**J-L**) ISH of *Cyp26b1* in the mitral valve between 12.5 and 18.5.

### *Cyp26b1* maintains aortic valve leaflet size by regulating cell proliferation

Due to its endocardial-specific expression in the cushions and heart valves, we asked if *Cyp26b1* is required for cardiac valve development and remodeling. To answer this question, we examined the heart valves at E18.5 in a *Cyp26b1* global knockout mouse (Daniel et al., 2020) (which normally die shortly after birth). In this mouse model, *Cyp26b1-/-* aortic valve leaflets at E18.5 appeared thickened compared to WT littermates (**Fig. 5A,B**). Quantification of valve leaflet widths revealed an approximate 2.5-fold increase in aortic valve leaflet width (**Fig. 5C**, n = 3 wildtype, 3 *Cyp26b1-/-* valves, *p* = 0.000049). This hypercellular increase was specific to the aortic valve, with the pulmonary, mitral, and tricuspid valve sizes being less affected by *Cyp26b1* deletion at E18.5. We asked if there were other cardiac defects in *Cyp26b1-/-* hearts, in addition to the aortic valve enlargement. Indeed, we observed a VSD (#, **Fig. 5E**) in 33% of *Cyp26b1-/-* hearts (n=1 out of 3) at E18.5 (**Fig. 5D-F**), indicating incomplete penetrance of a ventricular septal defect in embryos lacking *Cyp26b1.*

**Figure 5.**
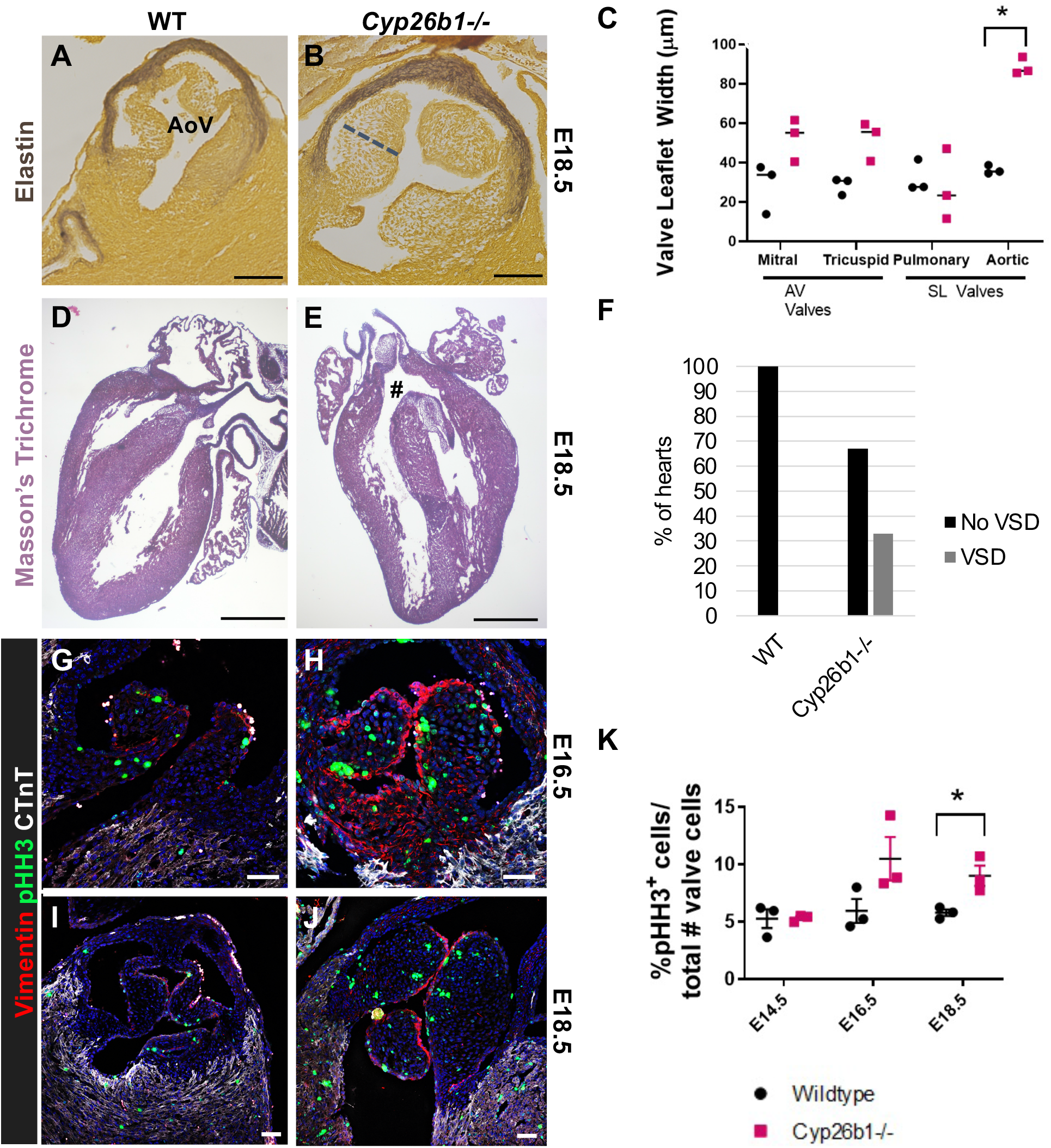
*Cyp26b1-/-* aortic valve leaflets are enlarged with aberrant cell proliferation. (**A,B**) Hart’s Elastin stain was performed on (**A**) WT and (**B**) *Cyp26b1-/-* embryonic heart sections at E18.5. (Scale = 100 μm). (**C**) The widths (μm) of WT (n=3) and *Cyp26b1-/-* (n=3) mitral, tricuspid, pulmonary, and aortic valve leaflets at E18.5 were quantified. (Statistical significance was carried out using multiple unpaired t-tests, with the Holm-Sidak method used to correct for multiple comparisons, * p<0.05.) (**D,E**) Masson’s Trichrome stain was performed on (**D**) WT and (**E**) *Cyp26b1-/-* embryonic heart sections at E18.5. (Scale = 500 μm). (**G-I**) Immunofluorescence analysis of paraffin sections of cardiac valve leaflets from WT and *Cyp26b1-/-* embryos at E16.5 and E18.5 was performed using antibodies against Vimentin, pHH3, and cardiac Troponin T. (Scale = 50 μm). (**J**) The percentage of pHH3+ cells per valve section was quantified (Statistical significance was carried out using multiple unpaired t-tests, with the Holm-Sidak method used to correct for multiple comparisons, * p<0.05.)

We then asked if there was increased cell proliferation specifically in *Cyp26b1-/-* aortic valve leaflets. Immunostaining was performed using anti-phospho-Histone H3 (pHH3) antibody, to ascertain the number of proliferating cells in the valves. Since the heart valves undergo normal extension and remodeling during the later stages of embryonic development (Lincoln et al., 2007), proliferating cells were expected to be seen in WT valves. Indeed, pHH3-expressing cells were present in WT valve leaflets at E16.5 and E18.5 (**Fig. 5G, I**). pHH3+ cells were also observed throughout the mesenchyme of both AV (**Fig. S5**) and SL valves from *Cyp26b1-/-* hearts at E16.5 and E18.5 **(Fig. 5G, H, J)**. The percentage of pHH3+ cells in each valve section was quantified, and no significant differences in proliferation were detected at E16.5 in either AV or SL valves. However, by E18.5 the proportion of pHH3+ cells was significantly increased from 5.7% in wildtype to 8.9% in *Cyp26b1-/-* aortic valve leaflets (**Fig. 5J**, n = 3 wildtype, 3 *Cyp26b1-/-* valves, p = 0.025). These data indicate that SL valve enlargement is due, at least in part, to increased cell proliferation in *Cyp26b1-/-* hearts, and that *Cyp26b1* is required to restrain cell proliferation during valve elongation and remodeling.

### *Cyp26b1* restrains RA targets in valve endothelial and mesenchymal cells

In the canonical RA pathway, retinoic acid is internalized into the cell and then binds its receptors RAR and RXR (Balmer and Blomhoff, 2002). RAR and RXR then translocate into the nucleus where they activate transcription of RA response genes. Because Cyp26b1 metabolizes retinoic acid, we asked whether RA signaling was affected by loss of Cyp26b1. To assess this possibility, we examined known targets of RA such as Pbx1, Hnf3α, and Ret that have shown to be retinoic acid-responsive in other endothelial contexts (Balmer and Blomhoff, 2002; Daniel et al., 2020; Oppenheimer et al., 2007; Qin et al., 2004). We performed immunofluorescence for these downstream RA targets. At E15.5, we see broad increases in the expression of Pbx1 in the valve, as well as the surrounding aortic root (**Fig. 6A-B, Figure S6A**). At E15.5, Ret was not detected in the wildtype valve endothelium, but was expressed in ECs of *Cyp26b1-/-* embryonic valves (**Fig. 6C-D, C’-D’**). At E15.5, Hnf3α showed a broad increase in the valve endothelium in *Cyp26b1-/-* hearts compared to WT littermates (**Fig. 6E-F**). By E18.5, Hnf3α continues to be increased in the underlying mesenchyme (**Fig. S6B**). Together, these data suggest retinoic acid target genes are upregulated in the *Cyp26b1-/-* embryos, indicating dysfunction in RA metabolism. We propose that Cyp26b1 acts to metabolize circulating retinoic acid near the SL valves, and that loss of Cyp26b1 induces retinoic acid signaling to promote dysregulated mesenchymal cell proliferation (**Fig. 6G**).

**Figure 6.**
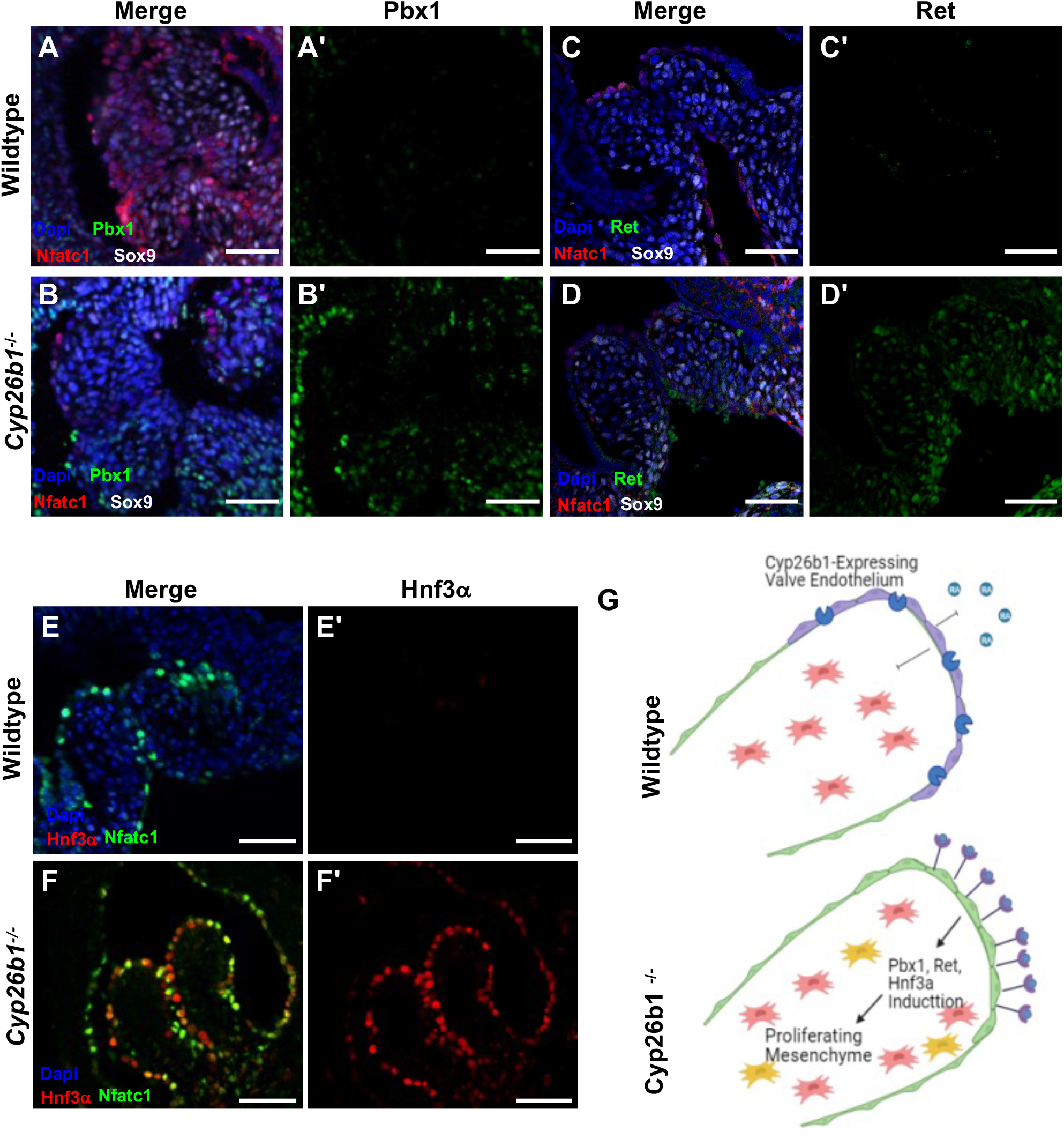
RA targets are elevated in the heart in the absence of Cyp26b1 by E15.5. Immunofluorescence for nfatc1 (heart valve endothelium), sox9 (mesenchymal cells), and retinoic acid target Pbx1 on paraffin sections of (**A**) Wildtype and (**B**) *Cyp26b1-/-* semilunar (SL) valves at E15.5. (**A’-B’**) Single channel images of Pbx1 immunofluorescence in wildtype (**A’**) and (**B’**) *Cyp26b1-/-* tissue. Immunofluorescence for nfatc1, sox9, and retinoic acid target Ret on paraffin sections of (C) Wildtype and (**D**) *Cyp26b1-/-* SL valves at E15.5. (**C’-D’**) Single channel images of Ret immunofluorescence in wildtype (**C’**) and (**D’**) *Cyp26b1-/-* tissue. Immunofluorescence for nfatc1, sox9, and retinoic acid target Hnf3α on paraffin sections of (**E**) Wildtype and (**F**) *Cyp26b1-/-* SL valves at E15.5. (**E’-F’**) Single channel images of Hnf3α immunofluorescence in wildtype (**E’**) and (**F’**) *Cyp26b1-/-* tissue. Scale bar represents 50μm. Images are representative results from three separate embryos**. (G**) Model of Cyp26b1 in valve development. In wildtype embryos, cyp26b1 metabolizes retinoic acid and restrains proliferation of the mesenchymal cells. In the absence of Cyp26b1, valvular endothelial cells experience an excess of RA signaling and induce expression of retinoic acid target genes. This leads an excess of mesenchymal cell proliferation and enlarged valve leaflets.

## DISCUSSION

Despite extensive clinical interest, endothelial contribution to cardiac valve development remains poorly understood, in part due to a dearth of markers for regionally restricted valve endothelial genes. In this report, we characterize dynamic endothelial gene expression within the developing heart valves. Furthermore, we identify *Cyp26b1* as enriched in both AV and SL valve endothelium and demonstrate its role during SL valve development. We show that Cyp26b1 is required to repress both endothelial and mesenchymal cell proliferation in the cardiac valves. In addition, we show that RA signaling targets are suppressed by Cyp26b1. Through these findings, our paper identifies a novel role for Cyp26b1 during heart valve elongation, highlights how regionally restricted genes contribute to valve development, and provides a useful set of tools to further probe endocardial heterogeneity. This study adds one more example of angiocrine signaling to a growing list of cross-signaling events known to occur in developing and adult tissues (Koch et al., 2021; Pasquier et al., 2020; Rafii et al., 2016).

The advent of single cell RNA sequencing has revealed a treasure trove of information, particularly endothelial data, that is ready for analysis by biologists. However, we suggest this technique has important shortcomings. The frequently limited amount of starting material used as input for single cell RNA seq can lead to high levels of uncertainty in the analysis (Lahnemann et al., 2020). Additionally, each single cell RNA seq experiment can generate several hundred to thousands of purportedly ‘differentially expressed’ candidate genes, that may or may not ultimately be validated, making it difficult to identify genes of significant interest. Here, we provide a method to prioritize candidate genes using publicly available databases, such as Genepaint and the Human Protein Atlas. We believe this methodology could and should be applied to verify single cell RNA seq data sets. As databases and datasets increase in numbers and become available, we propose it is imperative to cross-reference and validate bioinformatically generated data.

We found that published single cell RNAseq datasets do not always yield verifiable candidate genes that prove useful as endothelial markers. In one seminal study, ssRNAseq was performed on fetal heart valves ranging between 17- and 25-weeks post conception, a timeframe that corresponds to valve remodeling (Cui et al., 2019). However, we found that most genes identified were not expressed at levels high enough to detect upon validation with tools such as ISH or immunostaining (in our hands or in data available online). We acknowledge that there may be differences between mouse and human that account for these observations, however we have found similar validation discrepancies when analyzing mouse ssRNAseq datasets (data not shown). We demonstrate here that a pipeline of publicly available databases for gene expression mining, followed by traditional ISH, is useful for identifying useful markers (**Fig. S1**) and supports multimodal approaches. Genes displaying expression in both human and murine valve development are ripe for further functional follow-up.

Another advantage of creating a detailed and validated transcriptomic atlas of the developing heart is to deepen our understanding of the different molecular responses to the changing anatomy of the developing heart. During mid to late gestation in the mouse, ECs and underlying mesenchymal cells exhibit shifts in differential gene expression marking distinct domains. As valve mesenchyme diversifies during valve maturation, overlying endothelium displays distinct zones of gene expression, as assessed using both established and novel markers. *Lmo2* exhibits an expression pattern restricted to the inflow domain, while *Fzd10* and *Sgk3* show greater specificity for outflow domain expression. Identification of regionalized EC genes may help us answer open questions in the field. For instance, little is known about how valve cells transduce biomechanical signals stimulating the formation of stratified ECM in mature leaflets. ECs are in immediate proximity to blood flow and are thus poised to interpret mechanical cues from flow and communicate to the underlying mesenchyme. Identifying domain specific EC genes will allow generation of endothelial Cre lines that will help elucidate how distinct hemodynamic patterns present at the tip, outflow, and inflow influence valve elongation and remodeling. Another understudied question is determining how physiological distinctions in differing heart valves arise. We showed that *Sgk3* displays differential expression patterns in AV and SL valve leaflets. Understanding the function of regionalized EC genes in the heart may clarify the cellular and molecular bases for morphological differences seen between different types of valves.

Our transcriptomic analysis of the developing mouse cardiac valves led us to identify Cyp26b1 as expressed in the early heart and as a potentially important factor in valve development. Because Cyp26 enzymes facilitate degradation of RA and are required to restrict RA during early development (Otto et al., 2003; Pennimpede et al., 2010; White et al., 1997), we further tested a role for Cyp26b1 in heart development. Strikingly, loss of Cyp26b1 had a profoundly detrimental impact on heart formation, leading to abnormally enlarged heart valves and VSDs. We show that Cyp26b1 is required for proper formation of the mouse ventricular septum and OFT and that RA signaling is increased upon loss of Cyp26b1. The powerful effects of RA on embryogenesis, and particularly on cardiogenesis, have long been known (Lammer et al., 1985; Pan and Baker, 2007; Wilson et al., 1953). Exogenous RA treatment or inhibition of Cyp26 enzyme function in animal models induces cardiac defects that mimic human CHDs (Roberts et al., 2006; Yasui et al., 1995). Our data align with these results.

How and why Cyp26b1 is expressed in all four heart valves, but only has a profound effect on aortic valves, is still unclear. We surmise that the answer might be found in the differential cellular contribution to the valves. In mouse embryos lacking the retinaldehyde dehydrogenase-2 enzyme (Raldh2), cardiac neural crest cells (cNCCs) were abnormally localized in the OFT endocardial cushions (El Robrini et al., 2016). Proper RA signaling is required by other cell types in the OFT to stimulate its formation and patterning (Keyte and Hutson, 2012). These studies and others showed that RA is necessary for proper allocation of cNCCs to the endocardial cushions. It is possible that loss of Cyp26b1 led to dysregulated cNCC behavior. An intriguing possibility is the role of RA-mediated induction of apoptosis in OFT cushions. Others have shown that apoptosis is increased in OFT cushions of *RXRa-/-* mutant mouse hearts (Kubalak et al., 2002), indicating a requirement for RA mediation of cell death in the developing OFT. Work from the Epstein lab has shown that cNCCs are required to stimulate mesenchymal cell apoptosis during late-stage remodeling and valvular thinning in the OFT (Jain et al., 2011). Intriguingly, their mouse models of cNCC-deficient OFTs displayed dysmorphic and thickened SL valves, similar to the *Cyp26b1-/-* aortic valve leaflets. Since cell death is required for proper valve leaflet elongation and remodeling, and RA signaling induces apoptosis in the OFT, it is possible that loss of Cyp26b1 interferes with cNCC-mediated mesenchymal cell apoptosis and remodeling.

On the other hand, pulmonary valve elongation and remodeling were relatively unaffected by loss of *Cyp26b1* in our mouse model, supporting the idea that the aortic valve defects and VSD may be cNCC-independent. Studies of Cyp26a1 and Cyp26c1 function in the zebrafish heart showed that Cyp26a1/c1 loss-of-function disrupts ventricular cardiomyocyte polarity and cardiac structure (Rydeen and Waxman, 2016). However, these studies do not distinguish whether these effects are cell-autonomous or not. It will be interesting to investigate whether there are cell-autonomous myocardial defects n *Cyp26b1-/-* mouse hearts. Additionally, the Waxman lab showed that RA deficiency leads to a smaller second heart field (SHF) population and a restricted arterial pole in embryonic zebrafish hearts(Duong et al., 2021), indicating that proper RA signaling is required for SHF progenitor differentiation into cardiomyocytes and smooth muscle cells in the OFT. Further investigation into cell-autonomous myocardial defects in *Cyp26b1-/-* mouse hearts is warranted.

Another question that arises is why the *Cyp26b1-/-* mutants display VSDs in only 30% of cases. We propose this could be an indirectly caused by valve defects. The ventricular septum (VS) contains both muscular and membranous components. The muscular VS grows first, followed by the membranous VS. The membranous VS arises due to interactions between the muscular VS and the AV endocardial cushion cells (Icten and Tetik, 1996). Studies have shown that the membranous VS is derived from the AV endocardial cushion lineage, and that proper morphogenesis of the OFT cushions is required for proper alignment and movement of the AV cushions (Komatsu et al., 2007). It is possible that defective fusion of OFT and/or AV endocardial cushions may lead to the VSD observed in some *Cyp26b1-/-* embryonic hearts. Another possibility is that *Cyp26b1-/-* hearts may contain myocardial defects that our analyses did not reveal. We suggest that at the very least *Cyp26b1-/-* VSDs indicates defects in alignment or cardiac remodeling, indicating a likely role for *Cyp26b1* beyond the valves themselves.

Finally, while loss of the RA-metabolizing enzyme Cyp26b1 might be expected to lead to increased RA signaling in the early valves, it remains unclear whether Cyp26b1-/- defects are fully explained by RA. RA treatment of avian embryos has indeed been shown to interfere with endocardial cushion development. In avian embryos treated at HH stage 15 (50 – 55 hr postfertilization), RA-treated hearts showed a significant volume decrease of the AV cushions by HH24 (4 dpf) in contrast to our findings. RA treatment also changed hemodynamics (Bouman et al., 1998). By HH34 (8 dpf), the AV valves were abnormally enlarged, possibly due to the blood flow defects and/or altered ECM composition (Bouman et al., 1997; Yan and Sinning, 2001). Therefore, while RA initially restricted endocardial cushion growth at HH24, hemodynamic changes in RA-treated embryonic hearts promoted abnormal expansion of heart valves by HH34. These experiments underscore how controlled RA signaling is crucial for proper cardiac development. In the case of avian cardiac development, and potentially in murine hearts lacking Cyp26b1, abnormally enlarged valves may result from cell-extrinsic effects downstream of aberrant RA.

In summary, in this report we have identified an atlas of genes that display an array of expression patterns during valve development using publicly available databases. Given that endothelial factors drive heart valve development, we have validated a number of these genes using ISH. We further investigated one particularly EC-restricted gene in the cardiac valves, *Cyp26b1,* and identified a previously unknown role for Cyp26b1 during the late stages of cardiac valve development and in VSD formation. We show that this phenotype is likely caused by an increase in retinoic acid signaling in valve ECs and underlying mesenchymal cells, as the downstream RA targets Pbx1, Ret and Hnf3α all display increased expression in these tissues. Our study highlights how dysfunction of a regionalized gene in the endothelium can have broad impacts in cardiac development, leading to CHD.

## MATERIALS AND METHODS

### Ethics statement

All animal experiments were performed in accordance with the Guide for the Care and Use of Laboratory Animals and the Animal Welfare Act, as well as protocols approved by the University of Texas Southwestern Medical Center Institutional Animal Care and Use Committee. All animals were observed daily, and appropriate care was provided by the veterinary staff of the UTSW Animal Resource Center, which is fully accredited by the Association for Assessment and Accreditation of Laboratory Care, International (Unit Number 000673) and by the NIH Office of Laboratory Animal Welfare (Assurance Number D16-00296). Dams were euthanized via IACUC-approved humane methods, using carbon dioxide asphyxiation and secondary cervical dislocation.

### Mice

Embryonic day 8.75 (E8.75) - E18.5 embryos (*Mus musculus*) were dissected and fixed in 4% PFA/PBS for 3 hours at 4°C for section immunofluorescence (IF) or overnight (O/N) for RNA ISH. Tissue was then washed, dehydrated to 70% ethanol (EtOH), and stored at −20°C. CD1 mice (Charles River Laboratories, Houston, TX) were used for WT experimental analyses. Otherwise, *Cyp26b1-/-* (Daniel et al., 2020) mice were used for experiments herein. Genotypes were determined by polymerase chain reaction (PCR) after O/N digestion using DirectPCR (Tail) Lysis buffer (Viagen, Los Angeles, CA) per manufacturer’s instructions.

### In-silico screening

Novel candidate markers suggestive of valve specific expression were identified by analyzing over 1000 gene expression patterns obtained via in-situ hybridization of paraffin sections of E14.5 mouse embryos depicted in the Genepaint database (https://gp3.mpg.de/). We also selected genes known to be involved in valve development.

Analysis of single-cell RNA-seq was carried out using the R package Seurat (Satija et al., 2015). The UMI count matrix provided by the Tang group (accession ID GSE106118) was uploaded into R. Cells with greater than 1000 genes and 5000 transcripts were retained for downstream analysis, as in the original paper (Cui et al., 2019). Expression data was log transformed, and we identified 2000 highly variable genes using the Seurat package. These genes were used to perform principal component analysis (PCA) in R. Significant PCs were identified using the Jackstraw function in R, with the number of replicate samplings set to 100. Eleven principal components (From PC1 to PC11) were used to perform t-SNE analysis, with clustering parameter resolution set to 0.5. Cluster identification was performed by looking at expression of known marker genes in t-SNE plots (See Fig. S2 for more details).

### RNA probe generation

Digoxigenin-labeled probes were synthesized as previously described (Xu et al., 2011). Briefly, plasmids (GE Dharmacon, Lafayette, CO) were linearized using restriction enzymes listed in **Table S1**. Antisense DigoxigeninUTP-labeled RNA probes were synthesized at 37°C using RNA DIG labeling mix per manufacturer’s instructions (Roche) using RNA polymerase. After incubation with RQ1 DNase I (Promega, Madison, WI), RNA was purified using Micro Bio-Spin columns (BioRad, Hercules, CA).

### Section RNA in situ hybridization

RNA in situ hybridization (ISH) on paraffin sections was performed as previously described (Braitsch et al., 2019). Embryonic mouse heart sections were de-paraffinized and rehydrated before treatment with proteinase K. Following PBS washes, sections were placed in pre-hybridization solution pre-warmed to 65°C. RNA probes in hybridization buffer were applied to slides and then incubated O/N at 65°C. The next day, slides were washed, blocked, and incubated with anti-Digoxigenin antibody (Roche) at 4°C O/N. Subsequently, the slides were washed and incubated in BM purple (Roche) O/N at 37°C. Slides were then washed, refixed in 4% PFA, dehydrated, and then mounted with coverslips using Permount (Fisher Scientific, Hampton, NH). Images of section ISH hybridizations were obtained with an inverted microscope using 5x, 10x, and 20x objective lenses. Findings were categorized according to endothelial and mesenchymal domain specific expression patterns.

### Whole mount RNA in situ hybridization

RNA ISH on whole-mount mouse embryos was performed as previously described (Xu et al., 2011). Briefly, E8.75-E9.5 embryos were collected, and their yolk sac, amnion and neural tube were punctured to facilitate penetration of reagents. Embryos were fixed overnight in 4% PFA and subsequently dehydrated to 100% Methanol for storage at −20°C. Embryos were then rehydrated to filtered pH=7.5 0.1% PBST (Tween-20) on day of use, and permeabilized with 10ug/mL Proteinase K (Boehringer 161 519) in PBST for 15 minutes at 37°C. Fixation was stopped by washing in fresh 2mg/mL glycine in PBST for 5min. Following rinses in PBST, the embryos were re-fixed with 0.2% glutaraldehyde in 4% paraformaldehyde in PBST for 20 minutes at room temperature (RT). After rinsing embryos in PBST, embryos were incubated in pre-heated PreHyb solution (50% formamide (Fisher BP 227 100), 5x SSC pH=4.5, 50ug/mL ribonucleic acid from torula yeast, type VI (Sigma R6625), 1% SDS, 50ug/mL Heparin (Sigma H4784)) for 1 hour at 65°C. Cyp26b1 probe was then diluted in PreHyb buffer and embryos were incubated in probe overnight at 65°C.

The next day, embryos were washed with pre-warmed Solution I (50% formamide, 20% 20x SSC pH=4.5, 5% 20% SDS in water) two times at 65°C. Embryos were transitioned to Solution II (10% 5M NaCl, 1% 1M Tris pH=7.5, 0.1% Tween-20 in water, filtered) by washing in a 1:1 mix of Solution I and Solution II for 10 minutes at 65°C. Embryos were washed with Solution II for 10 minutes three times, and then washed in 100ug/mL RNase A (Sigma R 6513) in Solution II for 1 hour at 37°C. This was followed by one rinse in Solution II and one rinse in pre-warmed Solution III (50% formamide, 10% 20x SSC pH=4.5, 1% 20% SDS in water) at RT. Embryos were then washed with Solution III 2 times for 30 minutes at 65°C. Embryos were transitioned to MBST (100mM Maleic acid (Sigma M 0375), 150mM NaCl pH=7.5, 0.1% Tween-20 in water) for three rinses. Embryos were pre-blocked with 2% Boehringer Mannheim Blocking Reagent (BMB 1 096 176) for 3-4 hours at RT. Following blocking, embryos were incubated overnight at 4°C in 1:4,000 antibody mix (Anti-Digoxigenin-AP Fab Fragments (same as in situ hybridization on sectioned tissue)) diluted in blocking solution.

The next day, embryos were rinsed with MBST 3 times, and washed with MBST for 1hr 8 times at RT. Embryos were rinsed 3 times in NTMT (2% 5M NaCl, 5% 2M Tris pH=9.5, 5% 1M MgCl_2_, 0.1% Tween-20 in water) and then incubated in BM Purple (Boehringer Mannheim 1 442 074) for at least 1 hour. Once samples were dark enough with purple signal, embryos were rinsed 3 times in PBST pH=4.5 and then fixed in 0.1% glutaraldehyde in 4% paraformaldehyde in PBST for at least 1hr or overnight. After rinsing 3 times in PBST pH=4.5, embryos were visualized either in PBST or in 80% glycerol in PBST. Whole mount ISH images were taken with a NeoLumar stereomicroscope (Zeiss) using a DP-70 camera (Olympus).

### Immunofluorescence

Immunostaining was performed as previously described in (Braitsch et al., 2019). Tissue sections were deparaffinized, washed, and permeabilized using 0.3% Triton-X/PBS. Antigen retrieval was performed under pressure using acidic buffer A (Electron Microscopy Services, Hatfield, PA). Sections were then washed, blocked in CAS block (Invitrogen, Carlsbad, CA) and incubated in primary antibodies at 4°C overnight (for dilutions, see **Table S2**). Slides were then washed in PBS, incubated in Alexa Fluor secondary antibodies for 1-2 hours at RT, and subsequently incubated in 4’,6-diamidino-2-phenylindole dihydrochloride (DAPI) nuclear stain (Invitrogen). Slides were then washed in PBS and mounted using Prolong Gold Mounting Medium. Images were obtained using either a Nikon A1R or a LSM710 Meta Zeiss confocal microscope.

### Immunohistochemistry

H&E staining was performed as previously described (Daniel et al., 2020). Masson’s trichrome and Hart’s elastin staining were performed using standard protocols by the UTSW Pathology Core.

### Statistical Data

The width (μm) of WT (n=3) and *Cyp26b1-/-* (n=3) mitral, tricuspid, pulmonary, and aortic valve leaflets at E18.5 were measured using FIJI. (Statistical significance was carried out using multiple unpaired t-tests, with the Holm-Sidak method used to correct for multiple comparisons, * p<0.05.)

Proliferative (pHH3+/DAPI+/cTnT-) valve cells were assessed on sections of E14.5, E16.5, and E18.5 hearts. In WT and *Cyp26b1-/-* hearts, the number of pHH3+ cells in the valve leaflet(s) on each section were quantified and normalized to the total number of DAPI+ valve nuclei per section, and at least three sections were quantified per valve (n=3 embryos per stage per genotype). Data are presented as mean ± Standard Error of the Mean (SEM). Unpaired Student’s *t* test was performed to compare the percentage of proliferating cells, as well as valve leaflet width, by using GraphPad and Microsoft Excel. A *p* value <0.05 was considered to be statistically significant (*).

## ACKNOWLEDGEMENTS

We are grateful to Steven Lu from the STARS 2019 summer program for initial work on the Cyp26b1 mutant aortic valve enlargement and VSDs. We thank the UTSW Transgenic Core for initial generation of the Cyp26b1 mice, as per (Daniel et al., 2020). Thank you to Thomas Carroll for the Wnt4 plasmid. Thank you to Ethan Black, Thejal Anandakumar, Vedha Vaddaraju and Kate Black for help screening through hundreds of candidates online. We are grateful to the Cleaver lab for discussions and critique throughout this project. We are grateful to the SURISKD project for summer support of E.B.

## Author contributions

Conceptualization: C.M., M.H., N.A., O.C.; Methodology: C.M., M.H., N.A., O.C.; Validation: C.M., M.H., N.A., E.B., T.A., V.V., S.V.; Investigation: C.M., M.H., N.A., H.B.; Resources: C.M., M.H., N.A., O.C.; Data curation: C.M., M.H., N.A., E.B., T.A., V.V., O.C.; Writing – original draft: C.M., M.H., N.A., O.C.; Writing – review and editing: C.M., M.H., N.A., H.R.B., S.V., O.C.; Visualization: C.M., M.H., N.A.; Supervision: O.C.; Project administration: O.C.; Funding acquisition: O.C.

## Author contributions

This work was supported by the National Heart Lung and Blood institute (HL113498 to O.C.); the National Institute of Diabetes and Digestive and Kidney Diseases (DK106743, DK079862 to O.C.); and a Graduate Research Fellowship (2019241092 to H.R.B.). Deposited in PMC for release after 12 months.

## Supplementary information

Supplementary information available online at

