## Supplementary Materials (figs, tables) for "Cyp26b1 restrains murine heart valve growth during development"

### Supplementary Figures

#### SUPPLEMENTARY FIGURE LEGENDS

**Figure S1**. **Genes expressed in the cardiac valves of E14.5 mouse embryos.** (**A**) Transcriptomic analysis of genes using public databases yielded numerous genes expressed either throughout the AV and SL valve endothelium and mesenchyme, (**B**) primarily in the valve endothelium, (**C**) and throughout the valve endothelium as well as trabecular endocardium. All individual gene expression images are from Genepaint (https://gp3.mpg.de/).

**Figure S2**. **Single Seq Cluster Identification.** (**A**) *Myh7*, *Tnnt2*, *Tbx5,* and *Actn2* were used to identify cardiomyocyte clusters. (**B**) *Tbx18*, *Tcf21*, *Sema3d,* and *Wt1* (Xu et al., 2009; Zeng et al., 2011) were used to identify epicardial clusters. (**C**) *Sox9*, *Edil3*, *Eln,* and *Cd9* were used to identify valve mesenchyme clusters (Gitler et al., 2003; Hinton et al., 2010). **(D)** *Tpsab1, csf1r, cd7, and cd14* were used to identify immune and lymph cluster (Burel et al., 2019; Cui et al., 2019). (**E**) *Hbb* and *hbm* were used to identify erythrocytes (Cui et al., 2019).

**Figure S3. Known valve markers.** (**A-L**) RNA ISH using probes against known valve genes was performed on paraffin sections of embryonic mouse hearts at E10.5 and E14.5. (**A-C**) *Sox9* is robustly expressed in the cushion and valve mesenchyme (Lincoln et al., 2007). (**D-F**) *Wnt4* expression is endothelial-specific in the E10.5 cushions. At E14.5, *Wnt4* expression is highest in the tip and inflow regions of the AV and SL valves (Alfieri et al., 2010). (**G-I**) At E10.5, Car3 is expressed sporadically in the cushion mesenchyme (Maeda et al., 2019). In the AV valves at E14.5, *Car3* is most highly expressed in the endothelial inflow domain and ventricularis mesenchymal domain. In the SL valves *Car3* is concentrated in the endothelial inflow and mesenchymal atrialis domains**.** (**J-L**) *Ccn3* expression is concentrated within the tip and distal mesenchymal regions of the cushion at E10.5 (Lin et al., 2010). At E14.5, *Ccn3* shows endothelial and mesenchymal expression in the AV valves, while the SL valve has strong expression of *Ccn3* in the tip domain. Scale = 100 μm.

**Figure S4**. **At E8.75, Cyp26b1 is expressed in the developing vasculature, but not the endocardium.** (**A**) Though valve endothelial cells clustered very tightly together (**A'**), *Cyp26b1* was not very robust in any of the clusters. (**B**) At E8.75, *Cyp26b1* ISH showed expression in the developing vasculature (**B''**), but not the endocardium (**B'**).

**Figure S5**. **Loss of Cyp26b1 does not affect cell proliferation or valve leaflet width at E15.5.** (**A-D**) Immunofluorescence analysis of paraffin sections of cardiac valve leaflets from WT and *Cyp26b1-/-* embryos at E16.5 and E18.5 was performed using antibodies against Vimentin, pHH3, and cardiac Troponin T. (Scale = 50 μm). (**E**) The percentage of pHH3+ cells per valve section was quantified and found to not be statistically elevated in mutant valves. (**F**)Valve leaflet width was quantified on H&E stained paraffin sections across all valves from WT and *Cyp26b1-/-* embryos at E15.5. (Statistical significance was carried out using multiple unpaired t-tests, with the Holm-Sidak method used to correct for multiple comparisons, * $p<0.05$.).

**Figure S6**. **Retinoic acid target genes are expressed in throughout the aortic valve and aortic root.** Immunofluorescence for Nfatc1 (heart valve endothelium), Sox9 (mesenchymal cells), and retinoic acid target Pbx1 on paraffin sections of (**A**) Wildtype and (**B**) *Cyp26b1-/-* semilunar (SL) valves at E15.5. **(A'- B')** Single channel images of Pbx1 immunofluorescence in wildtype (**A'**) and (**B'**) *Cyp26b1-/-* tissue. (**C**) Immunofluorescence retinoic acid target Hnf3α on paraffin sections of

(**C**) Wildtype and (**D**) *Cyp26b1-/-* SL valves at E15.5. Scale bar represents 50 μm.  Images represent results of three independent experiments.

#### SUPPLEMENTARY INFORMATION REFERENCES


**Alfieri, C. M., Cheek, J., Chakraborty, S. and Yutzey, K. E.** (2010). Wnt signaling in heart valve development and osteogenic gene induction. *Dev Biol* **338**, 127-135.

**Burel, J. G., Pomaznoy, M., Lindestam Arlehamn, C. S., Weiskopf, D., da Silva Antunes, R., Jung, Y., Babor, M., Schulten, V., Seumois, G., Greenbaum, J. A., et al.** (2019). Circulating T cell-monocyte complexes are markers of immune perturbations. *Elife* **8**.

**Cui, Y., Zheng, Y., Liu, X., Yan, L., Fan, X., Yong, J., Hu, Y., Dong, J., Li, Q., Wu, X., et al.** (2019). Single-Cell Transcriptome Analysis Maps the Developmental Track of the Human Heart. *Cell Rep* **26**, 1934-1950 e1935.

**Gitler, A. D., Lu, M. M., Jiang, Y. Q., Epstein, J. A. and Gruber, P. J.** (2003). Molecular markers of cardiac endocardial cushion development. *Dev Dyn* **228**, 643-650.

**Hinton, R. B., Adelman-Brown, J., Witt, S., Krishnamurthy, V. K., Osinska, H., Sakthivel, B., James, J. F., Li, D. Y., Narmoneva, D. A., Mecham, R. P., et al.** (2010). Elastin haploinsufficiency results in progressive aortic valve malformation and latent valve disease in a mouse model. *Circ Res* **107**, 549-557.

**Lin, Z., Natesan, V., Shi, H., Hamik, A., Kawanami, D., Hao, C., Mahabaleshwar, G. H., Wang, W., Jin, Z. G., Atkins, G. B., et al.** (2010). A novel role of CCN3 in regulating endothelial inflammation. *J Cell Commun Signal* **4**, 141-153.

**Lincoln, J., Kist, R., Scherer, G. and Yutzey, K. E.** (2007). Sox9 is required for precursor cell expansion and extracellular matrix organization during mouse heart valve development. *Dev Biol* **305**, 120-132.

**Maeda, K., Ma, X., Hanley, F. L. and Riemer, R. K.** (2019). Modeling Impaired Coaptation Effects on Mitral Leaflet Homeostasis Using a Flow-Culture Bioreactor. *Ann Thorac Surg* **107**, 512-518.

**Xu, X. Q., Soo, S. Y., Sun, W. and Zweigerdt, R.** (2009). Global expression profile of highly enriched cardiomyocytes derived from human embryonic stem cells. *Stem cells (Dayton, Ohio)* **27**, 2163-2174.

**Zeng, B., Ren, X. F., Cao, F., Zhou, X. Y. and Zhang, J.** (2011). Developmental patterns and characteristics of epicardial cell markers Tbx18 and Wt1 in murine embryonic heart. *J Biomed Sci* **18**, 67.

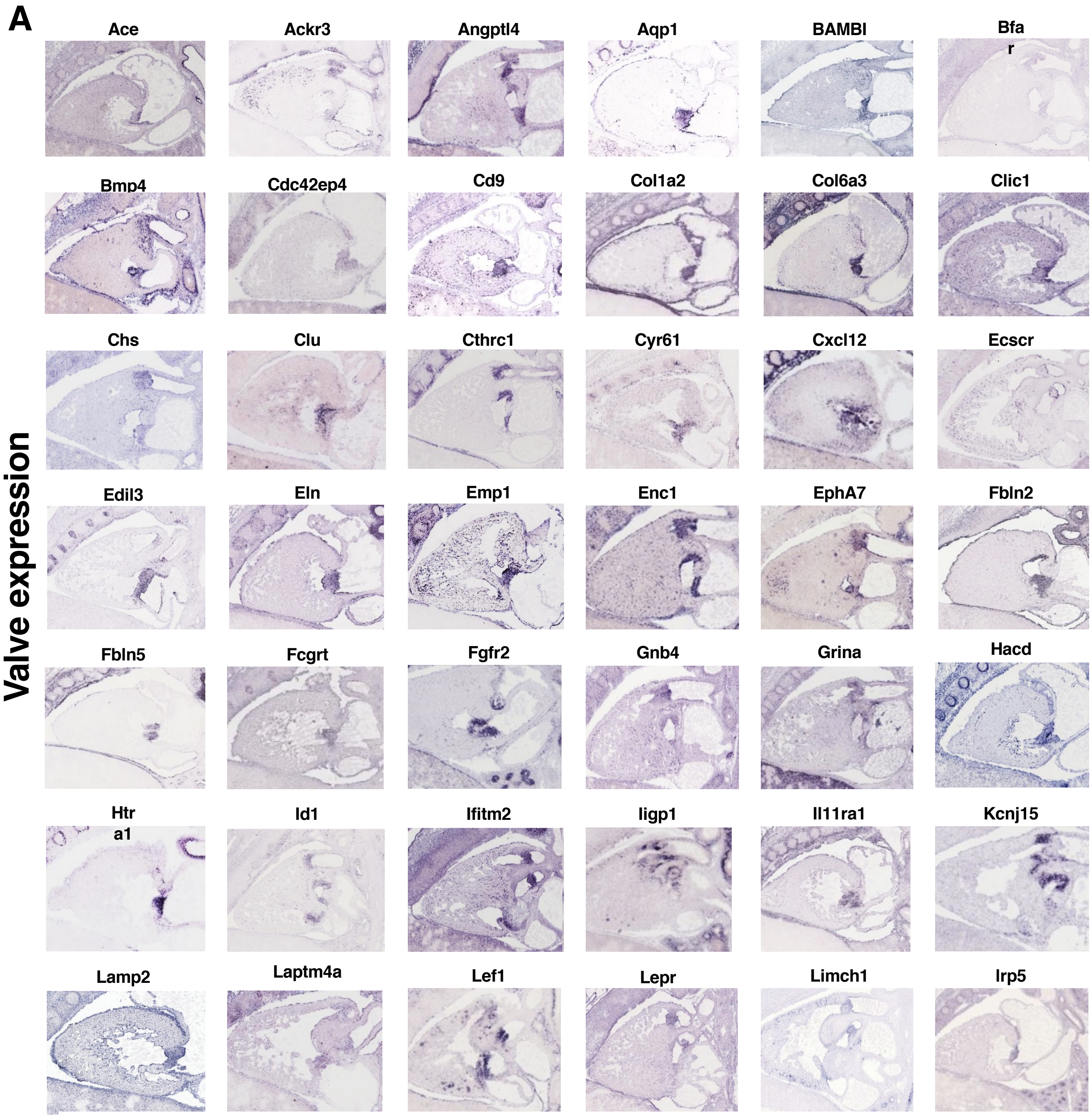


**Fig. S1. Expression patterns of candidates in heart valves at E14.5.**

A (Cont'd)

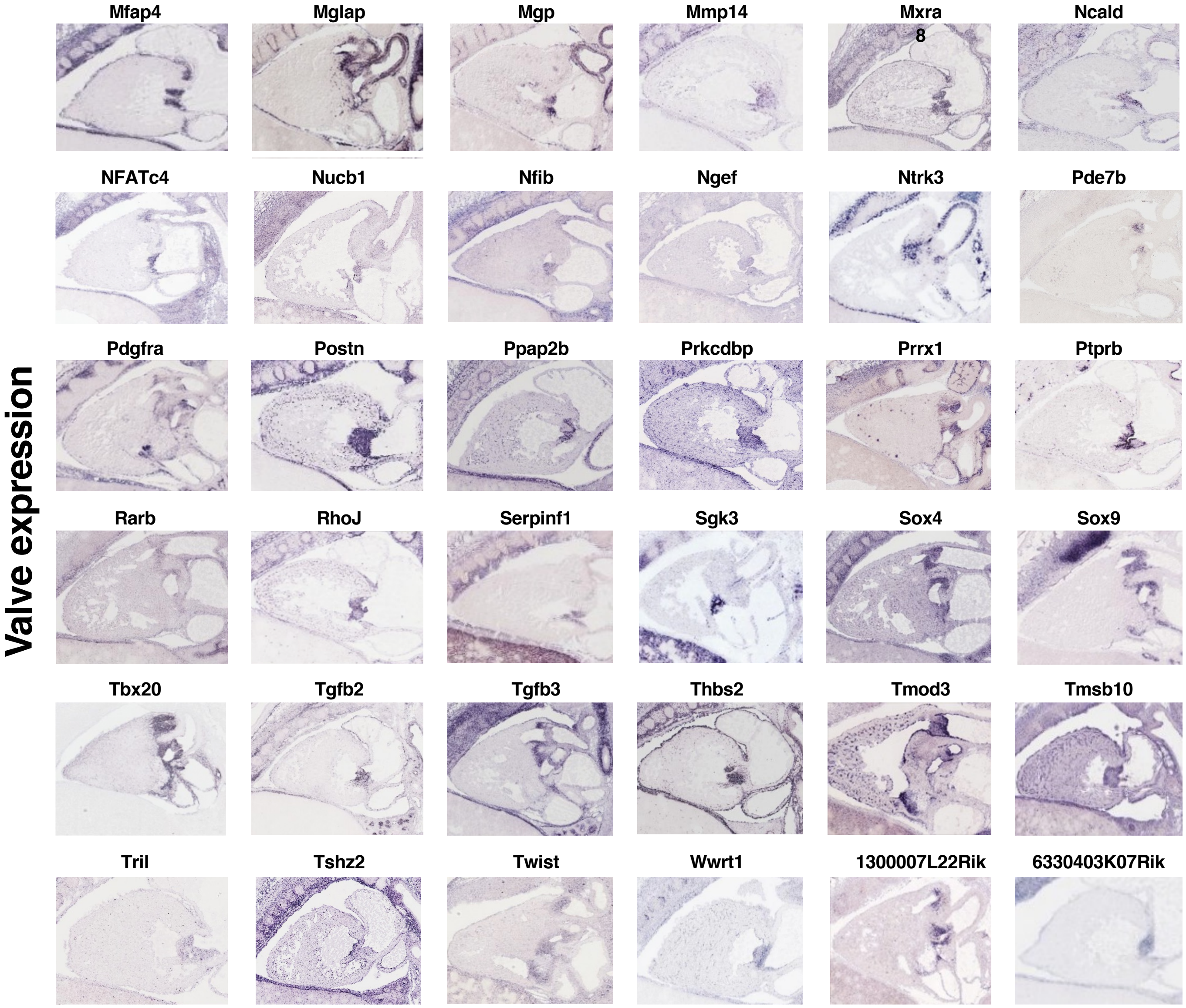


**Fig. S1. Expression patterns of candidates in heart valves at E14.5.**

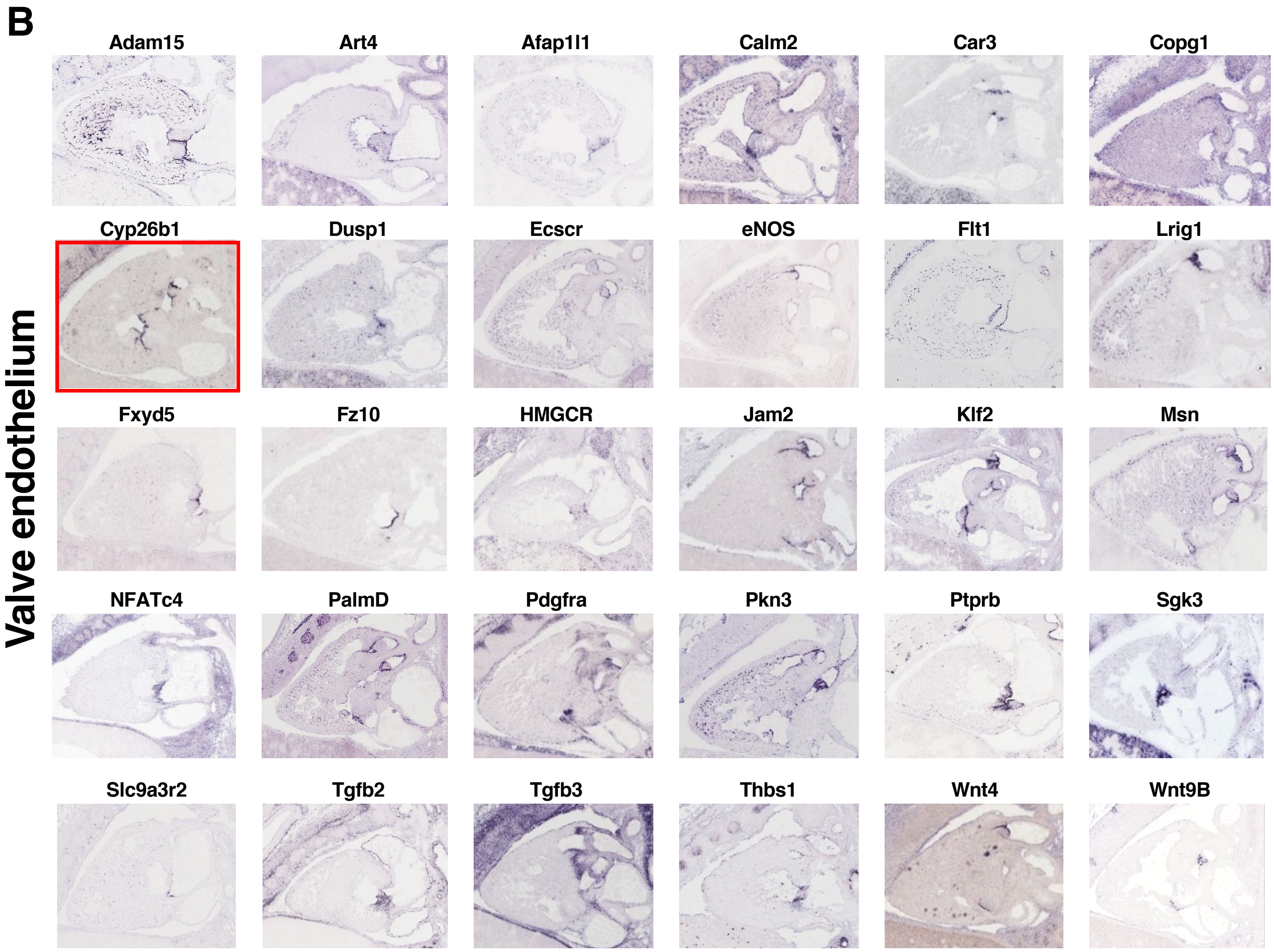


**Fig. S1. Expression patterns of candidates in heart valves at E14.5.**

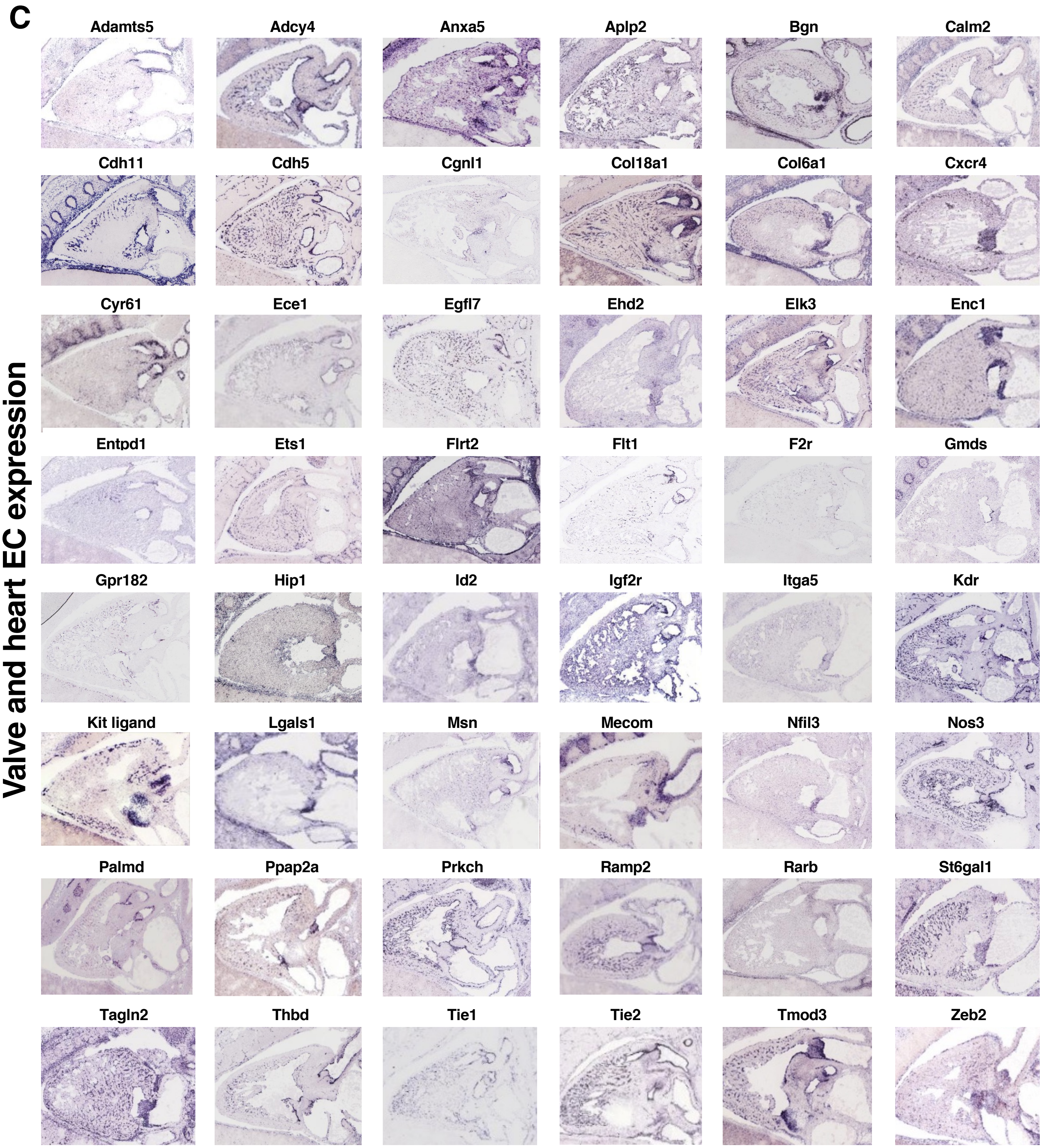


**Fig. S1. Expression patterns of candidates in heart valves at E14.5.**

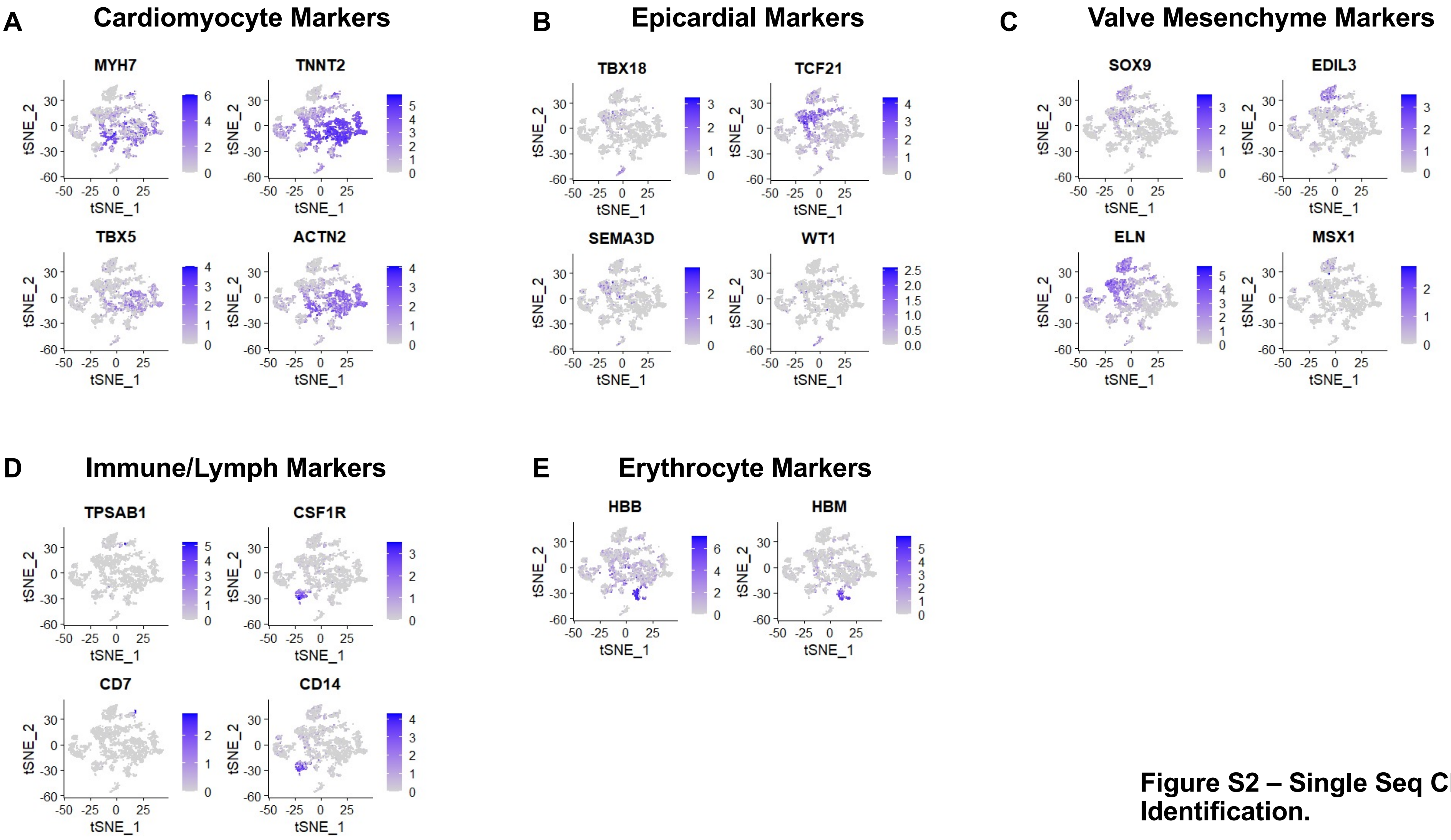


**Figure S2 – Single Seq Cluster Identification.**

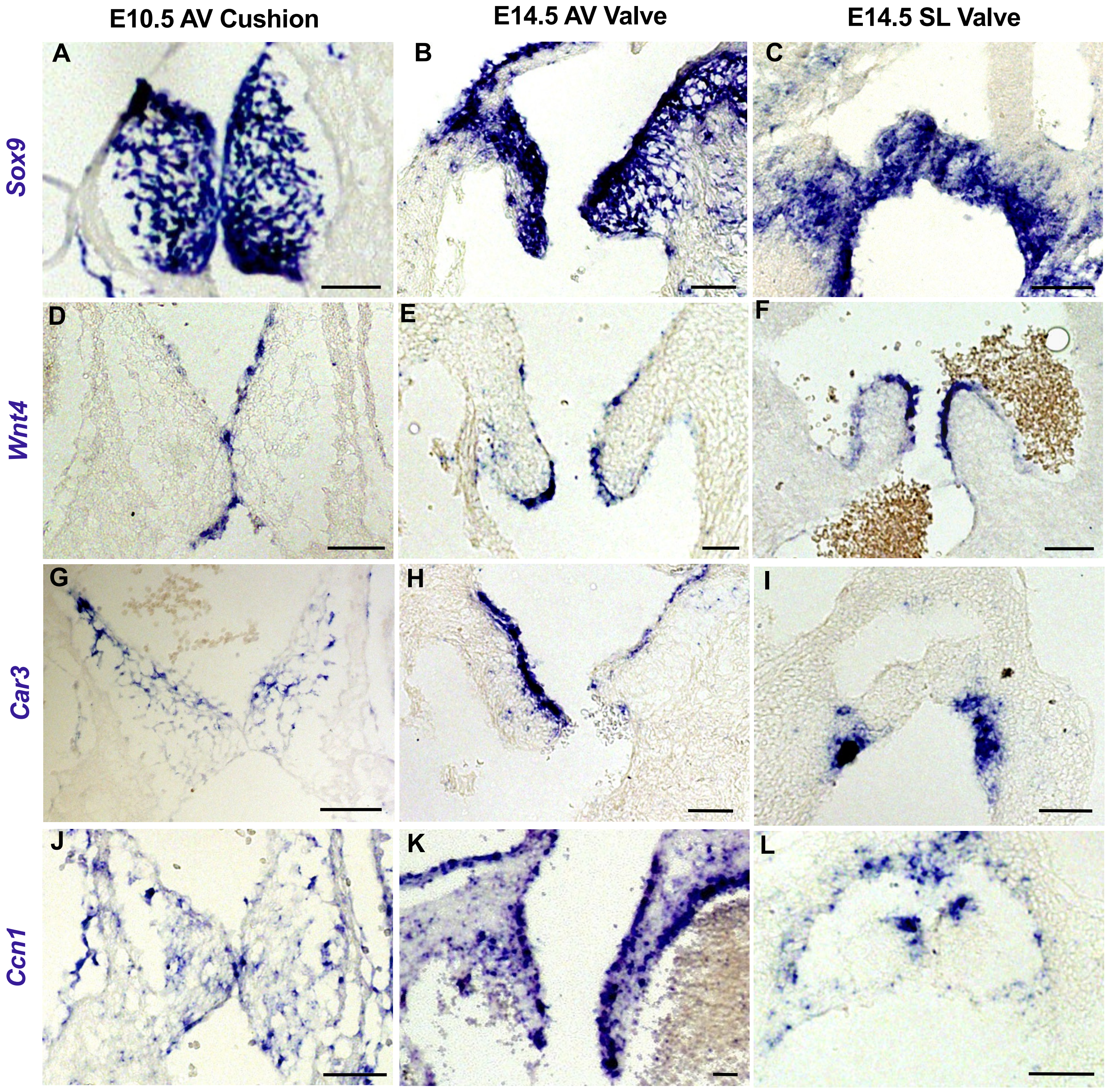


**Figure S3 – Known valve markers.**

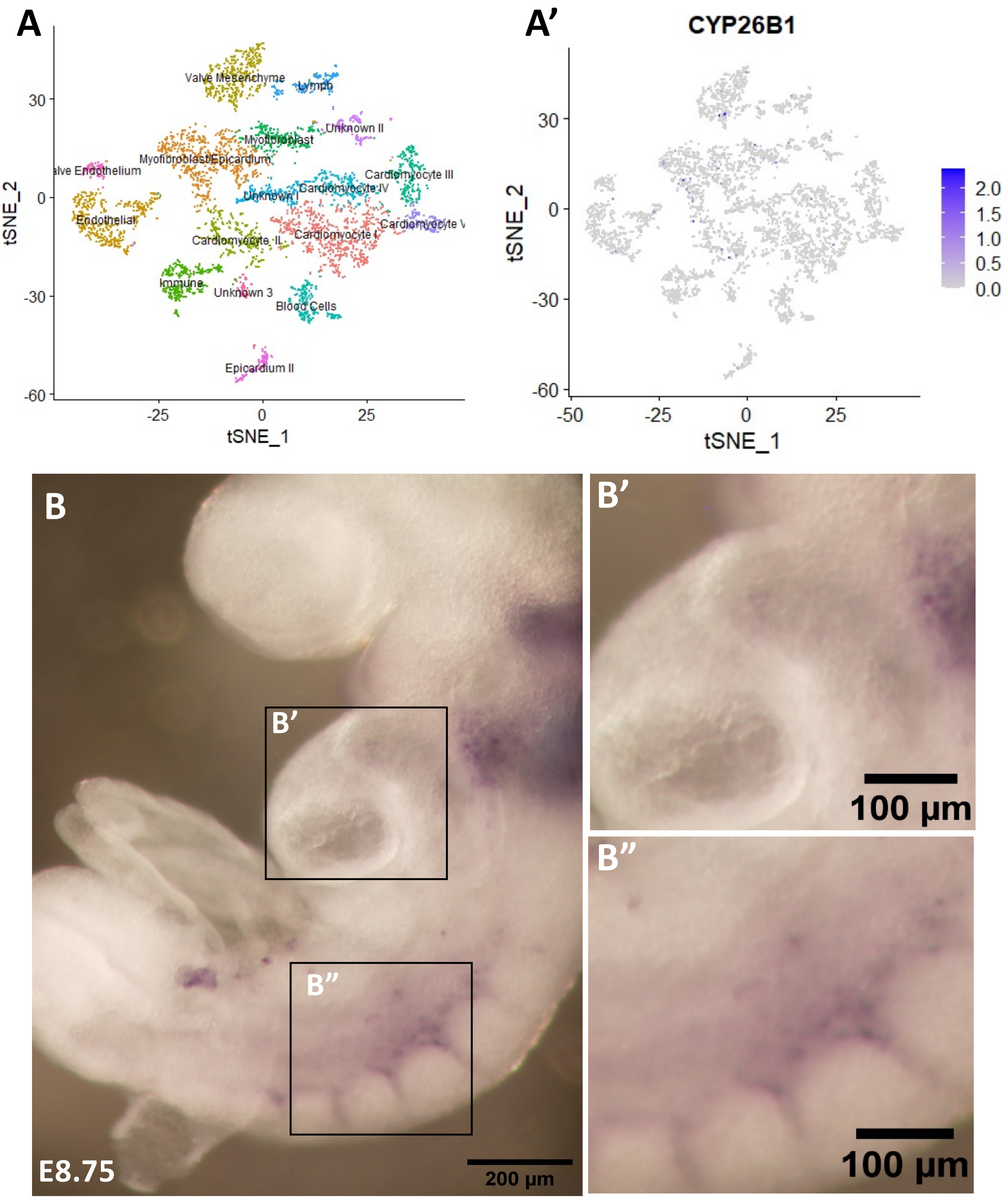


**Figure S4 – At E8.75, Cyp26b1 is expressed in the developing vasculature, but not the endocardium.**

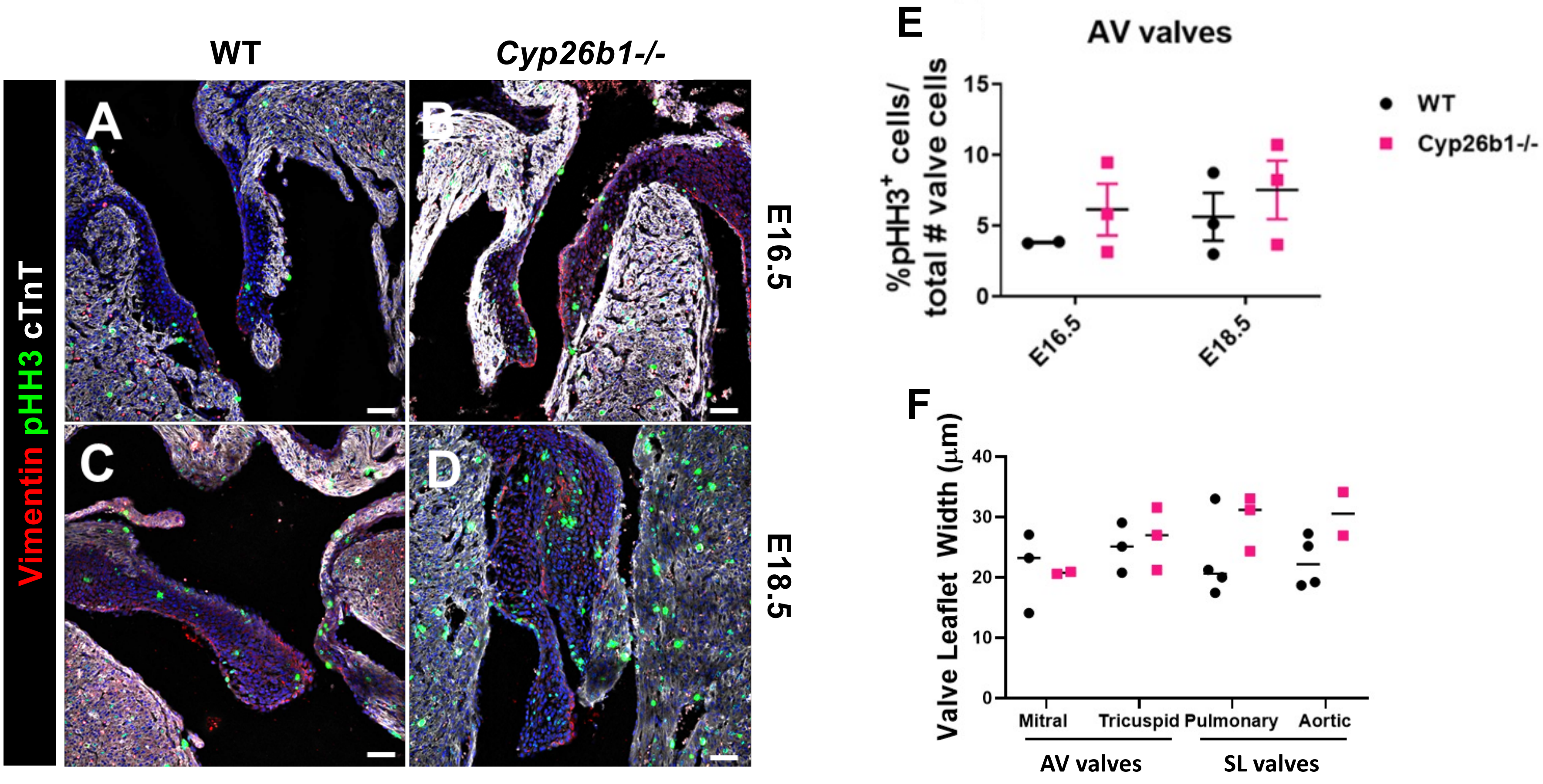


**Figure S5 – Loss of Cyp26b1 does not affect cell proliferation or valve leaflet width at E15.5.**

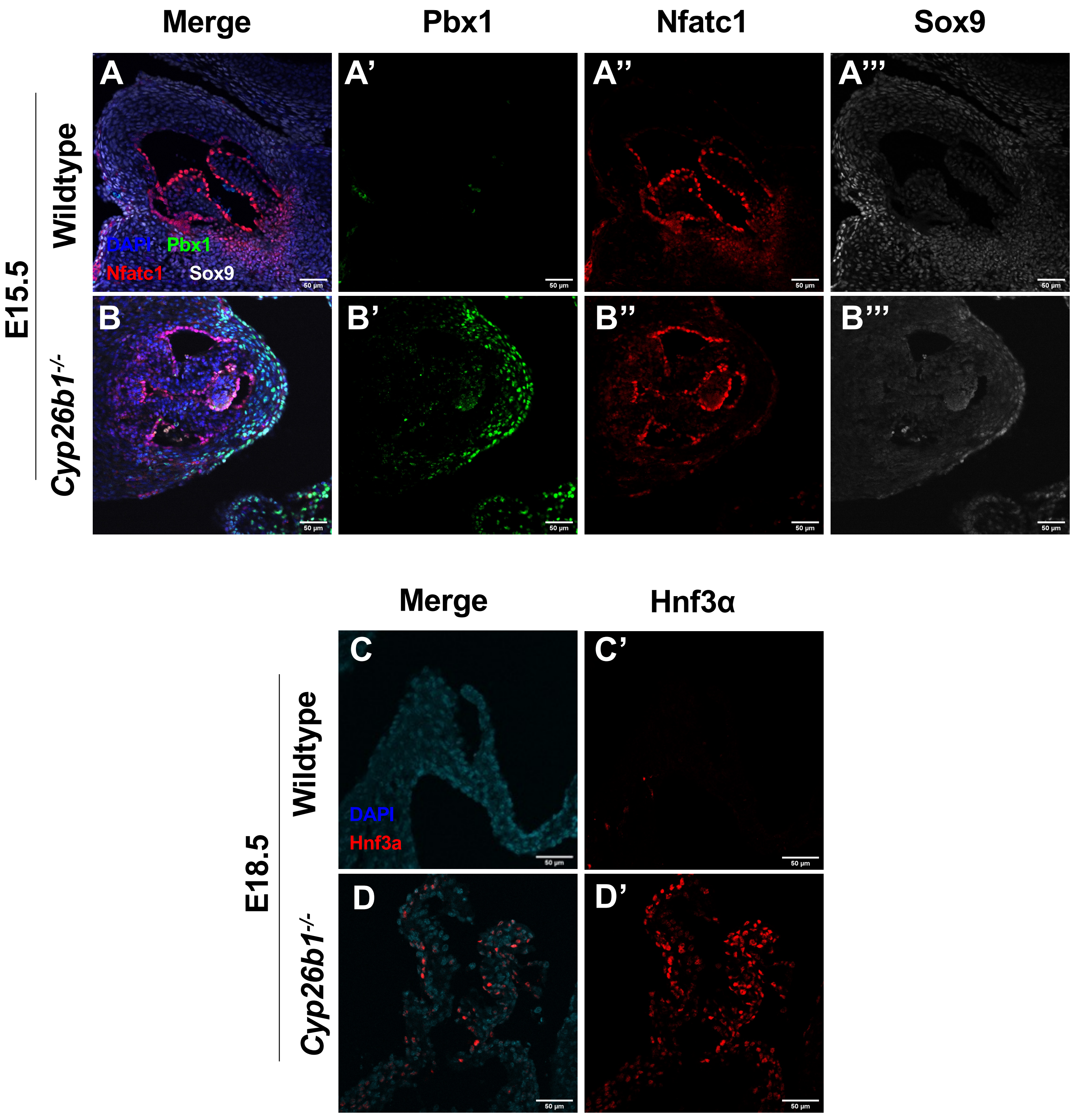


**Figure S6 – Retinoic acid target genes are expressed throughout the aortic valve and aortic root**

**S1 Table. Clones used for probe generation for in situ hybridization.**

| Clone | CloneID (Dharmacon) | Accession Number | Plasmid | Enzyme | RNA Polymerase |
|---|---|---|---|---|---|
| *Car3* | 4195712 | BC011129 | pCMV-SPORT6 | *Sal1* | T7 |
| *Cyp26b1* | 6400154 | BC059246 | pYX-Asc | *EcoRI* | T3 |
| *Cyr61* | 5716887 | BC066019 | pYX-Asc | *EcoRI* | T3 |
| *Enc1* | 6853919 | CD348649 | pYX-Asc | *EcoRI* | T3 |
| *Fzd10* | 40090826 | BC117759 | PCR-BluntII TOPO | *EcoRV* | SP6 |
| *Lmo2* | 5349568 | BC057880 | pCMV-SPORT6 | *Sal1* | T7 |
| *Sgk3* | 5355819 | BI687953 | pCMV-SPORT6 | *Sal1* | T7 |
| *Sox9* | 5320371 | BC023953 | pCMV-SPORT6 | *Sal1* | T7 |
| *Wnt4* | ? | ? | ? | *Eag1* | T7 |

**S2 Table. Antibodies used for IF.**

| Name | Host Species | Company | Catalog # | Paraffin sections |
|---|---|---|---|---|
| Cardiac Troponin T | Mouse | Invitrogen | MA5-12960 | 1:100 |
| Phospho-Histone H3 | Rabbit | Millipore | 06-570 | 1:100 |
| Hnf3a | Mouse | Santa Cruz Biotechnology | sc-101058 | 1:100 |
| Nfatc1 | Mouse | Santa Cruz Biotechnology | sc-7290 | 1:100 |
| Pbx1 | Rabbit | Cell Signaling Technologies | 4342S | 1:100 |
| Ret | Rabbit | Cell Signaling Technologies | 14556T | 1:100 |
| Sox9 | Rabbit | Millipore | AB5535 | 1:700 |
| Vimentin | Goat | Santa Cruz | sc-7557 | 1:100 |
